# TREM2 regulates purinergic receptor-mediated calcium signaling and motility in human iPSC-derived microglia

**DOI:** 10.1101/2021.08.24.457491

**Authors:** Amit Jairaman, Amanda McQuade, Alberto Granzotto, You Jung Kang, Jean Paul Chadarevian, Sunil Gandhi, Ian Parker, Ian Smith, Hansang Cho, Stefano L. Sensi, Shivashankar Othy, Mathew Blurton-Jones, Michael Cahalan

## Abstract

The membrane protein TREM2 (Triggering Receptor Expressed on Myeloid cells 2) regulates key microglial functions including phagocytosis and chemotaxis. Loss-of-function variants of TREM2 are associated with increased risk of Alzheimer’s disease (AD). Because abnormalities in Ca^2+^ signaling have been observed in several AD models, we investigated TREM2 regulation of Ca^2+^ signaling in human induced pluripotent stem cell-derived microglia (iPSC-microglia) with genetic deletion of TREM2. We found that iPSC-microglia lacking TREM2 (TREM2 KO) show exaggerated Ca^2+^ signals in response to purinergic agonists, such as ADP, that shape microglial injury responses. This ADP hypersensitivity, driven by increased expression of P2Y_12_ and P2Y_13_ receptors, results in greater release of Ca^2+^ from the endoplasmic reticulum (ER) stores, which triggers sustained Ca^2+^ influx through Orai channels and alters cell motility in TREM2 KO microglia. Using iPSC-microglia expressing the genetically encoded Ca^2+^ probe, Salsa6f, we found that cytosolic Ca^2+^ tunes motility to a greater extent in TREM2 KO microglia. Despite showing greater overall displacement, TREM2 KO microglia exhibit reduced directional chemotaxis along ADP gradients. Accordingly, the chemotactic defect in TREM2 KO microglia was rescued by reducing cytosolic Ca^2+^ using a P2Y_12_ receptor antagonist. Our results show that loss of TREM2 confers a defect in microglial Ca^2+^ response to purinergic signals, suggesting a window of Ca^2+^ signaling for optimal microglial motility.

## Introduction

As the primary immune cells of the central nervous system, microglia survey their local environment to maintain homeostasis and respond to local brain injury or abnormal neuronal activity. Microglia are strongly implicated in several neurodevelopmental and neurodegenerative diseases^1–7^, warranting further study of human microglial dynamics. Purinergic metabolites (ATP, ADP, UTP, UDP) in the brain constitute key signals driving microglial activation and chemotaxis, and are detected by microglial cells over concentrations ranging from hundreds of nM to mM ^8–13^. ATP released from both homeostatic and damaged cells is hydrolyzed locally by nucleosidases such as the ectonucleotidase NTPDase1 (CD39) or pyrophosphatase NPP1 to produce ADP^14–16^. ADP is then detected by P2Y purinergic receptors on microglia, causing IP_3_-dependent Ca^2+^ release from the endoplasmic reticulum (ER) lumen. Ca^2+^ depletion from the ER in turn activates ER STIM1 proteins to translocate proximally to puncta where closely apposed plasma membrane (PM) Orai1 channels are activated. This mechanism underlies store-operated Ca^2+^ entry (SOCE) in many cell types ^17^, including microglia^18–20^.

Purinergic signaling is central to microglial communication with other brain cell types and has been negatively correlated with the onset of Disease-Associated Microglia (DAM) transcriptional states ^21–25^. P2Y_12_ and P2Y_13_ receptors are highly expressed by microglia and are activated predominantly by ADP ^16, 26^. P2Y_12_ receptors are essential for microglial chemotaxis and have been implicated in the microglial response to cortical injury^12, 27^, NLRP3 inflammasome activation^28, 29^, neuronal hyperactivity and protection^27, 30^, and blood brain barrier maintenance^31, 32^. While purinergic receptors have been broadly identified as markers of microglial homeostasis^23, 26^, mechanisms by which receptor expression may drive or maintain homeostatic microglial states remain incompletely understood.

Neuroinflammatory pathologies are often associated with altered Ca^2+^ signaling^33^. Microglia, in particular, show altered Ca^2+^ responses in mouse models of Alzheimer’s Disease (AD) by mechanisms that are not fully understood^34–36^. Ca^2+^ responses to purinergic metabolites have been extensively studied in cultured murine microglia, acute brain slices, and more recently in anesthetized mice ^8, 10, 34, 37–39^. However, our understanding of how specific patterns of Ca^2+^ signals in microglia correlate with and tune downstream microglial responses such as cell motility or process extension remains incomplete. There is also a paucity of knowledge on how regulators of purinergic Ca^2+^ signals in microglia might play a role in the dysregulation of Ca^2+^ signaling associated with aging and neuroinflammation.

TREM2 encodes a cell surface receptor that binds a variety of ligands including various lipids, apolipoprotein E (ApoE), and amyloid-β peptides. Upon ligand binding, TREM2 signals through its adaptor protein DAP12 to activate a host of downstream pathways^23, 40–42^. Loss of TREM2 function is thought to promote a more homeostatic-like state^23, 43, 44^. Indeed, microglia lacking TREM2 expression exhibit greatly diminished activation against disease pathology, correlating with increased risk of AD^23, 41, 45^. Purinergic receptor hyperexpression has been reported at the transcriptome level across multiple TREM2 loss of function models including human patient mutations ^21–23, 25, 41, 46^. For example, P2Y_12_ receptor protein expression was found to be elevated in the cortical microglia of *Trem2^-/-^* mice and in a preclinical mouse model of AD^47, 48^, although the mechanistic link between purinergic receptor expression and TREM2 function remains poorly understood.

We previously developed methods to generate human induced pluripotent stem cell-derived microglia (iPSC-microglia)^49–51^, which can be used to model human microglial behavior. While iPSC-microglia are proving increasingly useful to investigate neurodegenerative disorders^41, 52–56^, Ca^2+^ signaling has not yet been extensively profiled in these models. In this study, we compared purinergic Ca^2+^ signaling and motility characteristics in WT and TREM2 KO human iPSC-microglia, and examined the mechanisms that underlie enhanced purinergic Ca^2+^ signaling in microglia lacking TREM2. We find that motility is differentially tuned by Ca^2+^ in TREM2 KO cells with consequences for chemotaxis.

## Results

### Purinergic receptor Ca^2+^ signaling is enhanced in TREM2-knockout human iPSC-microglia

To determine if TREM2 plays a role in microglial Ca^2+^ signaling, we compared cytosolic Ca^2+^ responses to the purinergic agonist ADP in isogenic, CRISPR-modified wild type (WT) and TREM2-knockout (TREM2 KO) human iPSC-microglia. ADP stimulation induced a biphasic Ca^2+^ response – a rapid initial peak followed by a secondary phase of sustained Ca^2+^ elevation lasting several minutes, in line with previous observations in mouse microglia ^57, 58^. Both phases of the Ca^2+^ response were significantly elevated in TREM2 KO microglia, raising the possibility that augmentation of the initial Ca^2+^ response to ADP in TREM2 KO microglia may be coupled to a larger sustained component of Ca^2+^ entry (Figure 1A, B). These results were corroborated in iPSC-derived microglia cell line expressing the genetically-encoded Ca^2+^ indicator Salsa6f^59, 60^ (Figure 1C, D). The Salsa6f probe showed the expected increase in the GCaMP6f fluorescence in response to Ca^2+^ elevation without any change in the tdTomato signal, and it did not perturb microglial activation and function (Figure 1-figure supplement 1A-G). TREM2 KO microglia also showed exaggerated Ca^2+^ responses to the purinergic agonists ATP and UTP at similar low mM concentrations, although the secondary Ca^2+^ elevations were not as long-lasting as with ADP (Figure 1E, F **and** Figure 1-figure supplement 2).

**Figure 1:**
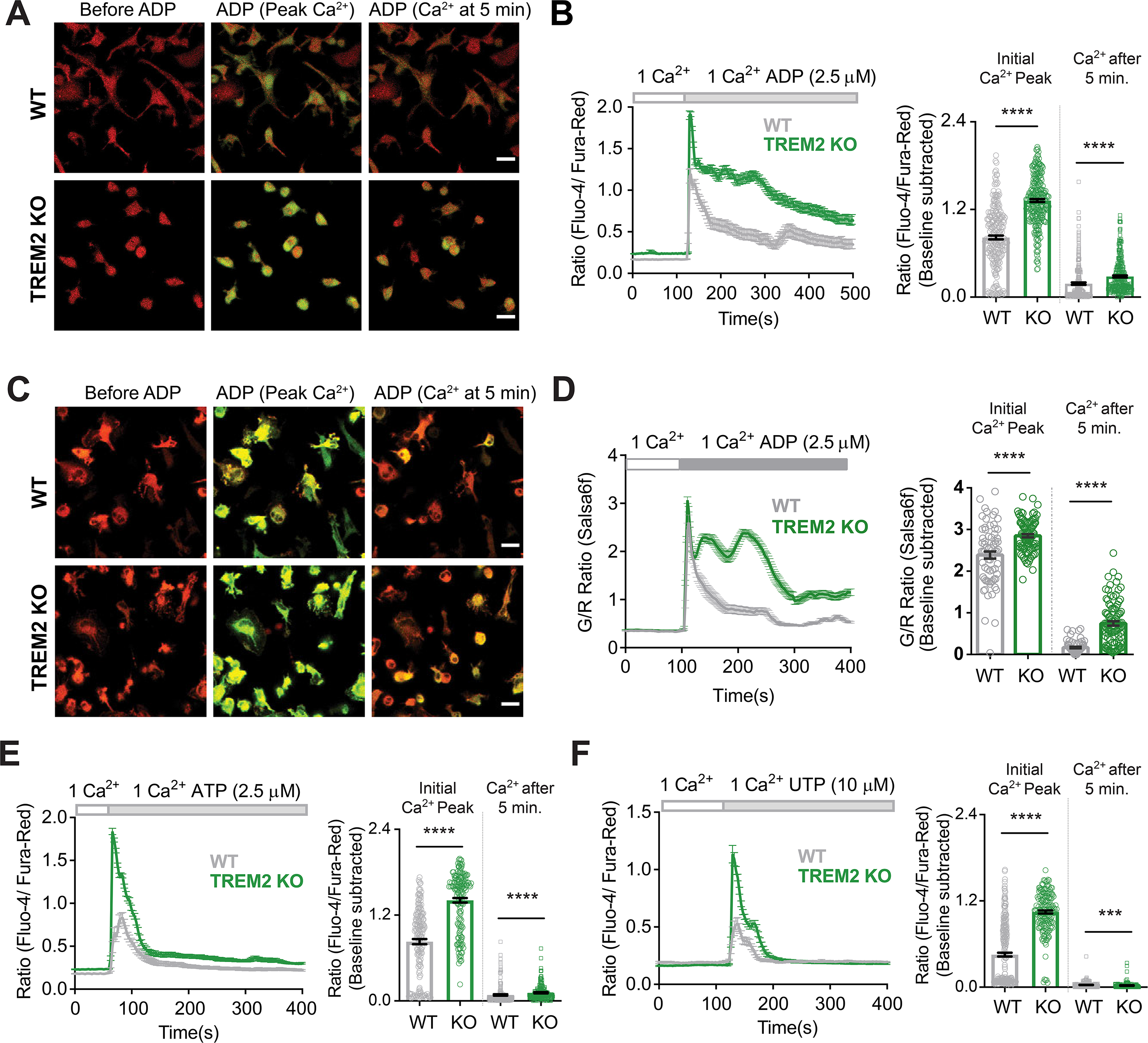
Microglia lacking TREM2 show exaggerated Ca^2+^ responses to purinergic stimulation. (**A**) Representative red-green channel overlay images of WT (top) and TREM2 KO (bottom) iPSC-microglia loaded with Fluo-4 (green) and Fura-red (red) showing resting cytosolic Ca^2+^ before ADP, and Ca^2+^ levels 15 sec and 5 min after ADP addition. Scale bar= 20 μm. (**B)** Average traces (left panels) showing changes in cytosolic Ca^2+^ in response to 2.5 μM ADP in 1 mM Ca^2+^ buffer (n=39-44 cells). Baseline-subtracted peak Ca^2+^ response and cytosolic Ca^2+^ levels 5 min after ADP shown on the right (n=250-274 cells, 5 experiments, Mann-Whitney test). (**C-D**) Cytosolic Ca^2+^ response to ADP as in **A** and **B** but in iPSC microglia expressing the GCaMP6f-tdTomato fusion Ca^2+^ probe Salsa6f (n=41-53 cells, 2 independent experiments, Mann-Whitney test). Images in (**C**) are overlay of GCaMP6f (green) and tdTomato (red) channel images. Scale bar= 20 mm. (**E**) Ca^2+^ responses to 2.5 μM ATP in WT and TREM2 KO iPSC-microglia. Average traces (left panel, n=63-71 cells) and bar-graph summary of peak cytosolic Ca^2+^ and Ca^2+^ after 5 min (right panel, 165-179 cells, 3 experiments, Mann-Whitney test). (**F**) Ca^2+^ responses to 10 μM UTP. Average traces (45-55 cells) and summary of peak cytosolic Ca^2+^ and Ca^2+^ after 5 min (175-269 cells, 3 experiments, Mann-Whitney test). Data shown as mean ± SEM for traces and bar-graphs. *P values* indicated by *** for *P < 0.001,* **** for *P < 0.0001*.

### Increased P2Y_12_ and P2Y_13_ receptor expression drives increased peak Ca^2+^ in TREM2-KO microglia

Given the critical importance of ADP signaling in several aspects of microglial function, we investigated the mechanisms driving higher ADP-evoked Ca^2+^ signals in TREM2 KO microglia by focusing on specific steps in the purinergic Ca^2+^ signaling pathway (Figure 2A). The initial Ca^2+^ response to P2Y receptor engagement results from G protein-coupled phospholipase C activation, and IP_3_-mediated ER Ca^2+^ store-release. To test this, we treated cells with ADP in Ca^2+^-free solution buffered with the Ca^2+^ chelator EGTA to isolate Ca^2+^ signals from store-release and eliminate Ca^2+^ influx across the PM. Both WT and TREM2 KO cells exhibited a single Ca^2+^ peak, with TREM2 KO cells showing significantly higher peak Ca^2+^ response to ADP (Figure 2B **and** Figure 2-figure supplement 1A, B). Moreover, the amplitude of the Ca^2+^ peak was not significantly different in the presence or absence of external Ca^2+^, strongly suggesting that it is driven primarily by release of Ca^2+^ from intracellular stores even when external Ca^2+^ is present (Figure 2-figure supplement 1C). Dose-response curves for the peak Ca^2+^ response showed a steep leftward shift in TREM2 KO cells (Figure 2C). The EC_50_ value for WT microglia was 650 nM, whereas TREM2 KO microglia reached their EC_50_ by 15 nM. This stark difference was driven at least in part by a diminished percentage of WT cells responding to ADP at low μM doses (Figure 2D). However, limiting the analysis to cells that showed a Ca^2+^ rise revealed that “responding” TREM2 KO cells still exhibited higher Ca^2+^ responses to ADP than “responding” WT cells (Figure 2E). TREM2 KO microglia are thus significantly more sensitive to ADP than WT cells which may be critical in sensing ADP and in detecting ADP gradients.

**Figure 2:**
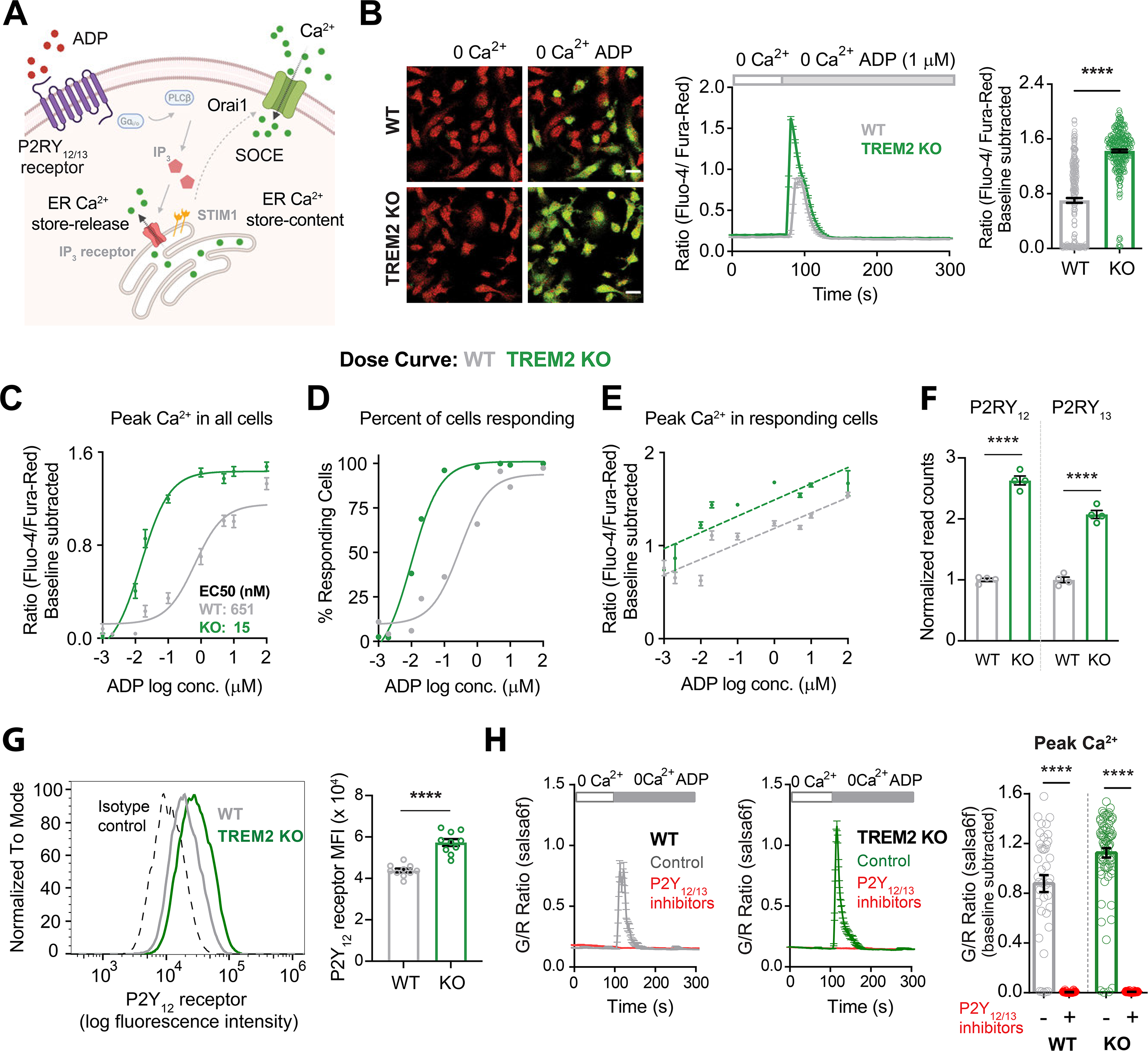
Higher sensitivity of TREM2 KO microglia to ADP is driven by increased purinergic receptor expression. (**A**) Schematic highlighting key downstream Ca^2+^ signaling events triggered by ADP. Cytosolic Ca^2+^ response to ADP is determined by functional expression and activity of P2Y_12_ and P2Y_13_ receptors, IP_3_ receptors, ER-store Ca^2+^ content, and store-operated Ca^2+^ entry (SOCE) regulated by STIM and Orai proteins. (**B**) Representative images (left panel) showing overlay of Fluo-4 (green) and Fura-red (red) channels in WT (top) and TREM2 KO (bottom) iPSC-microglia before and peak Ca^2+^ response after ADP addition in Ca^2+^ free buffer. Scale bar= 20 μm. Average trace showing Ca^2+^ response to ADP in Ca^2+^-free buffer (middle panel, 64-83 cells). Quantification of peak signal (right panel, n=264-289 cells, 4 experiments, Mann-Whitney test). (**C-E**) Dose-response curves showing baseline-subtracted peak Ca^2+^ responses to ADP in Ca^2+^-free buffer (**C**), percent of “responding” cells (**D**) and peak Ca^2+^ responses only in “responding” cells (**E**). N= 84-474 WT cells and 70-468 TREM2 KO cells, 2-5 experiments. (**F**) RNA normalized read counts of P2Y_12_ and P2Y_13_ receptor expression from bulk RNA-sequencing of WT and TREM2 KO iPSC-microglia (n=4, adjusted *P-values* from DESeq2). (**G**) Representative histogram (left panel) showing PM expression of P2Y_12_ receptor in WT and TREM2 KO microglia. Cells were stained with BV421 labelled anti-human P2Y_12_ receptor antibody. Isotype control is shown as dashed line. Right panel shows summary of median fluorescence intensity (MFI) of P2Y_12_ receptor-labelled cells (n= 10 samples each, Students t-test). (**H**) Ca^2+^ traces (left panel) showing response to 1 mM ADP in Ca^2+^-free buffer after 30 min pre-treatment with a combination of P2Y_12_ receptor antagonist PSB 0739 (10 mM) and P2Y_13_ receptor antagonist MRS 2211 (10 mM). Summary of the peak Ca^2+^ response (right panel, n=40-79 cells, 2 experiments, Mann-Whitney test). Data are mean ± SEM. *P values* indicated by **** for *P < 0.0001*.

RNA-sequencing revealed significantly increased transcripts for P2Y_12_ and P2Y_13_ receptors, the main P2Y receptor subtypes in microglia that bind ADP, in TREM2 KO microglia^49, 50^ (Figure 2F). In comparison, relative mRNA levels of common mediators of Ca^2+^ signaling – including predominant isoforms of IP_3_ receptors, SOCE mediators Orai and STIM proteins, and SERCA and PMCA Ca^2+^ pumps – were either similar or modestly reduced in TREM2 KO in comparison with WT iPSC-microglia (Figure 2-figure supplement 1D, E). We therefore considered the possibility that signal amplification in microglia lacking TREM2 results primarily from increased expression of P2Y_12_ and P2Y_13_ receptors. Consistent with this, expression of P2Y_12_ receptors in the plasma membrane was significantly increased in TREM2 KO cells (Figure 2G). Furthermore, Ca^2+^ responses to ADP in Ca^2+^-free medium were completely abolished following treatment with a combination of P2Y_12_ and P2Y_13_ receptor antagonists (PSB 0739 and MRS 2211 respectively) in both WT and TREM2 KO microglia (Figure 2H). Treatment of cells with P2Y_12_ and P2Y_13_ receptor antagonists separately produced partial inhibition of peak ADP-mediated Ca^2+^ signals, implicating involvement of both receptor subtypes (Figure 2-figure supplement 1F, G). In summary, deletion of TREM2 results in a larger cytosolic Ca^2+^ peak in response to ADP due to increased expression of P2Y_12_ and P2Y_13_ receptors.

### SOCE through Orai channels mediates the sustained phase of ADP-evoked Ca^2+^ elevation

To probe the basis for the increased sustained component of ADP-evoked Ca^2+^ signal in TREM2 KO microglia, we examined SOCE using pharmacological and genetic approaches. Synta66, a reasonably specific inhibitor of Orai channels, significantly reduced the rate of SOCE following Ca^2+^ re-addition after ER store-depletion by the sarco-endoplasmic reticulum Ca^2+^ ATPase (SERCA pump) inhibitor, thapsigargin (TG) in both WT and TREM2 KO microglia (Figure 3A **and** Figure 3-figure supplement 1A). Using a similar Ca^2+^ re-addition protocol with ADP, we found significant inhibition of ADP-induced SOCE by Synta66 in both WT and TREM2 KO cells (Figure 3B **and** Figure 3-figure supplement 1B). The ADP-evoked sustained Ca^2+^ phase in TREM2 KO iPSC-microglia was also blocked by less specific Orai channel inhibitors, Gd^3+^ and 2-APB (Figure 3-figure supplement 1C, D). To further confirm the specific role of Orai1 channels in mediating SOCE, we generated an Orai1 CRISPR-knockout iPSC line. Deletion of Orai1 abrogated SOCE and significantly reduced the sustained Ca^2+^ response to ADP (Figure 3-figure supplement 1E, F). These results confirm that Orai1 plays an important role in mediating SOCE and ADP-evoked Ca^2+^ signals in iPSC-microglia.

**Figure 3:**
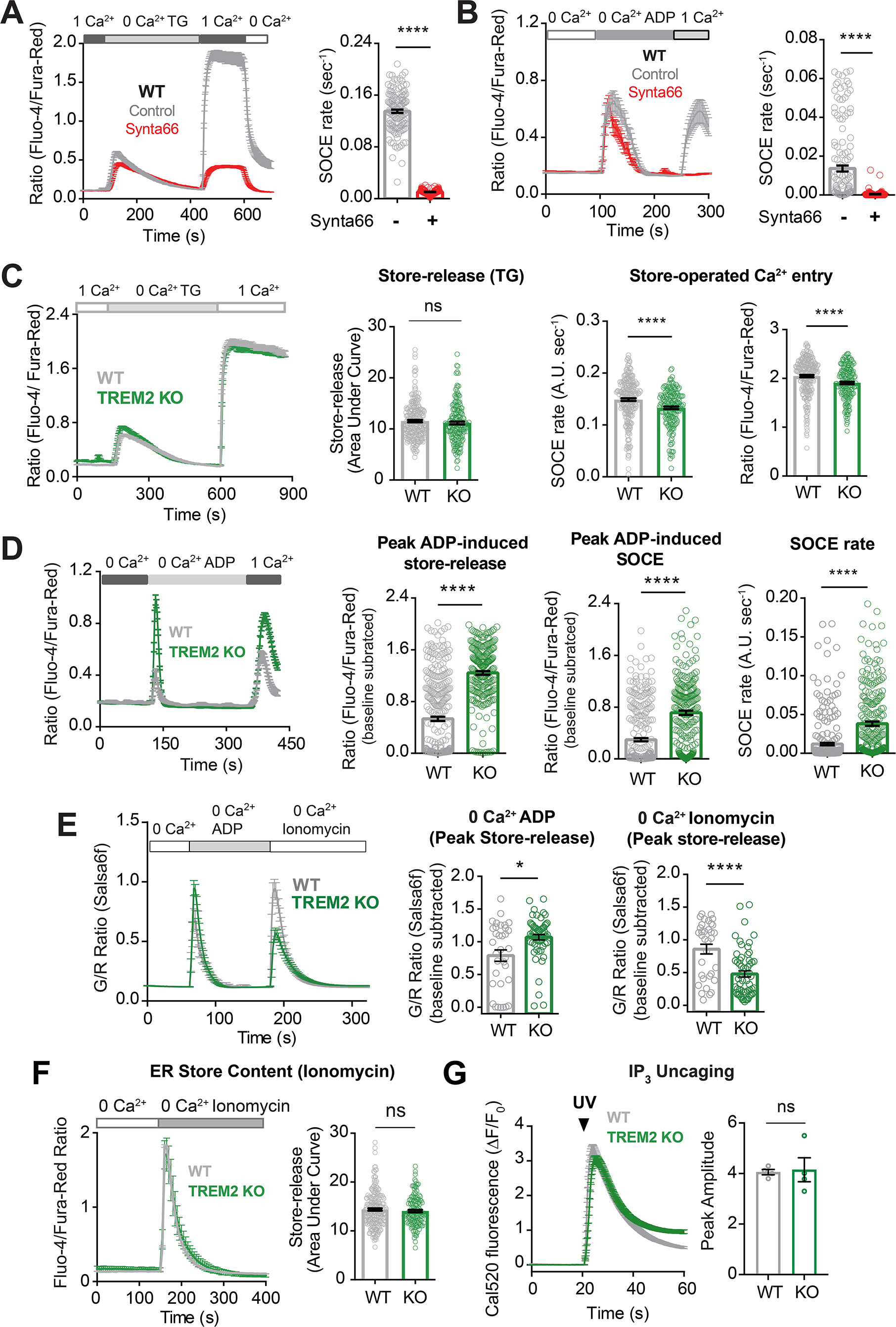
Regulation of ADP-evoked SOCE in WT and TREM2 KO microglia. (**A**) SOCE in WT microglia triggered with thapsigargin (TG, 2 μM) in Ca^2+^-free buffer followed by re-addition of 1 mM Ca^2+^ in the absence (control, grey trace) or presence (red trace) of the Orai channel inhibitor Synta66 (n=34-48 cells). Cells were pretreated with Synta66 (10 μM) for 30 min before imaging. Bar-graph summary of the rate of Ca^2+^ influx (n=80-137 cells, 2 experiments, Mann-Whitney test). (**B**) SOCE evoked by ADP (2.5 μM) in WT microglia (grey trace), using a similar Ca^2+^ addback protocol as in **A**. Red trace shows effect of Synta66 on ADP-evoked SOCE. Right panel shows bar-graph summary of the rate of ADP-triggered Ca^2+^ influx after re-addition of 1 mM Ca^2+^ (n=148-155 cells, 2 experiments, Mann-Whitney test). (**C**) Comparison of SOCE evoked with TG (2 μM) in WT and TREM2 KO cells (n=90-129 cells). Bar-graph summaries of ER store-release quantified as area under the curve, rate of SOCE, and peak SOCE (n=187-266 cells, 2 experiments, Mann-Whitney test). (**D**) Traces showing ADP-evoked SOCE in WT and TREM2 KO microglia after depleting stores with 100 nM ADP in Ca^2+^ free buffer and re-addition of 1 mM Ca^2+^ (left panel, n=97-114 cells). Comparison of ADP-evoked cytosolic Ca^2+^ peak, peak SOCE and SOCE rate (right panel, n=234-313 cells, 3 experiments, Mann-Whitney test). **(E)** Ionomycin-pulse experiment to measure residual ER Ca^2+^ pool in cells after initial treatment with ADP. WT and TREM2 KO cells were pulsed sequentially with ADP first (200 nM) and subsequently treated with ionomycin (1 μM) to empty and measure the residual pool of ER Ca^2+^. Imaging was done entirely in Ca^2+^-free buffer to prevent Ca^2+^ influx across the PM. Average trace (left panel), Peak ADP Ca^2+^ response (middle panel) and peak ionomycin-induced Ca^2+^ response (right panel) (n=38-60 cells, 3-4 experiments, Mann-Whitney test). **(F)** Average trace (left, 71-117 cells) and summary of ER store-release after 2 μM ionomycin treatment in Ca^2+^ free buffer (right, 146-234 cells, 2 experiments; **ns,** *non-significant P > 0.05,* Mann-Whitney test). (**G**) Same as H but in response to UV IP_3_ uncaging (167-200 cells, **ns***, non-significant P > 0.05,* nonparametric t-test). Data shown as mean ± SEM for traces and bar-graphs. Data are mean ± SEM. *P values* indicated by **ns** for non-significant, * for *P < 0.05* and **** for *P < 0.0001*.

To determine if SOCE is increased in TREM2 KO microglia and contributing to the higher sustained Ca^2+^ response to ADP, we compared the rate of store-operated Ca^2+^ influx after store-depletion with TG, and found that both the rate and amplitude of SOCE were modestly reduced in TREM2 KO cells (Figure 3C). In keeping with this, RNA sequencing revealed a modest reduction in STIM1 mRNA expression in TREM2 KO cells, although Orai1 mRNA was similar in WT and TREM2 KO microglia (Figure 2-figure supplement 1C, D). We further conclude that the elevated secondary phase of ADP-driven Ca^2+^ signals in TREM2 KO microglia is not primarily due to differences in the expression of STIM and Orai.

### ADP depletes ER Ca^2+^ stores to a greater extent in TREM2 KO microglia leading to greater SOCE activation

We hypothesized that the exaggerated secondary Ca^2+^ phase in response to ADP in TREM2 KO microglia may be driven by increased ER Ca^2+^ store-release leading to greater SOCE activation. Consistent with this possibility, peak cytosolic Ca^2+^ in response to partial store-depletion with ADP, and after Ca^2+^ re-addition was elevated in TREM2 KO microglia (Figure 3D). To examine if the higher magnitude of SOCE in TREM2 KO cells is due to depletion of ER Ca^2+^ stores by ADP, we sequentially treated cells with ADP followed by ionomycin to completely release stores in Ca^2+^ free buffer. While TREM2 KO cells showed greater peak Ca^2+^ with ADP as expected, the ionomycin Ca^2+^ peak – which reflects the residual ER Ca^2+^ pool – was significantly reduced indicating that ADP depletes ER Ca^2+^ stores to a greater extent in TREM2 KO cells (Figure 3E). Similar results were obtained when residual ER store-content was depleted using TG instead of ionomycin (Figure3-figure supplement 2A, B). We plotted cytosolic Ca^2+^ levels 5 minutes after addition of varying doses of ADP to indicate the degree of SOCE, as a function of the initial peak Ca^2+^, a readout of ER store-release (Figure 3-figure supplement 2C, D). Both WT and TREM2 KO microglia showed similar linear relationships between SOCE and store-release, further suggesting that SOCE is activated by similar mechanisms in the two cell lines, but is recruited to a greater extent in TREM2 KO cells due to increased ER store-release. We also note that increased sustained Ca^2+^ in TREM2 KO cells is unlikely to be due to differences in Ca^2+^ pump activity based on similar Ca^2+^ clearance rates (Figure 3-figure supplement 2E, F), consistent with comparable transcriptomic expression of major SERCA and plasma membrane Ca^2+^ ATPase (PMCA) isoforms in WT and TREM2 KO cells (Figure 2-figure supplement 1C, D).

Finally, quantification of cumulative cytosolic Ca^2+^ increases after maximally depleting ER stores with ionomycin alone suggested that overall ER store-content is not altered in microglia lacking TREM2 (Figure 3F). Comparison of Ca^2+^ responses to IP_3_ uncaging also ruled out major differences in the pool of functional IP_3_ receptors between WT and TREM2 KO cells (Figure 3G), as further substantiated by similar transcriptomic expression of IP_3_ receptor type 2 (the major IP_3_R subtype expressed in iPSC-microglia) in WT and TREM2 KO cells (Figure 2-figure supplement 1C, D)^41, 49^. In summary, deletion of TREM2 in iPSC-derived microglia leads to upregulation of P2Y_12_ and P2Y_13_ receptors and renders the cells hypersensitive to ADP signaling, consequently leading to greater IP_3_-mediated ER store-depletion and increased coupling to SOCE in response to purinergic metabolites.

### ADP potentiates cell motility and process extension in human WT iPSC-microglia

ADP is a potent chemoattractant for microglia^10^. Analogous to a previous study in fibroblasts^61^, we found that ADP treatment alters cell motility and leads to increased rates of scratch wound closure in WT iPSC-microglia (Figure 4A). To investigate the cellular mechanism of accelerated wound closure, we used time-lapse imaging to track open-field microglial cell motility (Figure 4B). Mean cell track speed and track displacement (defined as the overall change in position from the origin at a given time) were both increased after application of ADP. On the other hand, average track straightness, an indicator of how frequently cells change direction, was unaltered by ADP (Figure 4C). These data suggest that ADP-driven changes in motility in WT iPSC-microglia primarily arise from increases in microglial speed, and not altered turning behavior. ADP-dependent increases in speed were reversed in the presence of P2Y_12_ (PSB 0739) and P2Y_13_ (MRS 2211) receptor antagonists, confirming the role of these two purinergic receptors in ADP enhancement of microglial motility (Figure 4D). To determine if Ca^2+^ influx regulates ADP-mediated increases in motility, we measured cell migration with ADP in Ca^2+^-free medium and found that removing extracellular Ca^2+^ significantly decreased cell speed, displacement, and track straightness, suggesting that sustained Ca^2+^ signals are required for maximal increase in motility in response to ADP (Figure 4E).

**Figure 4:**
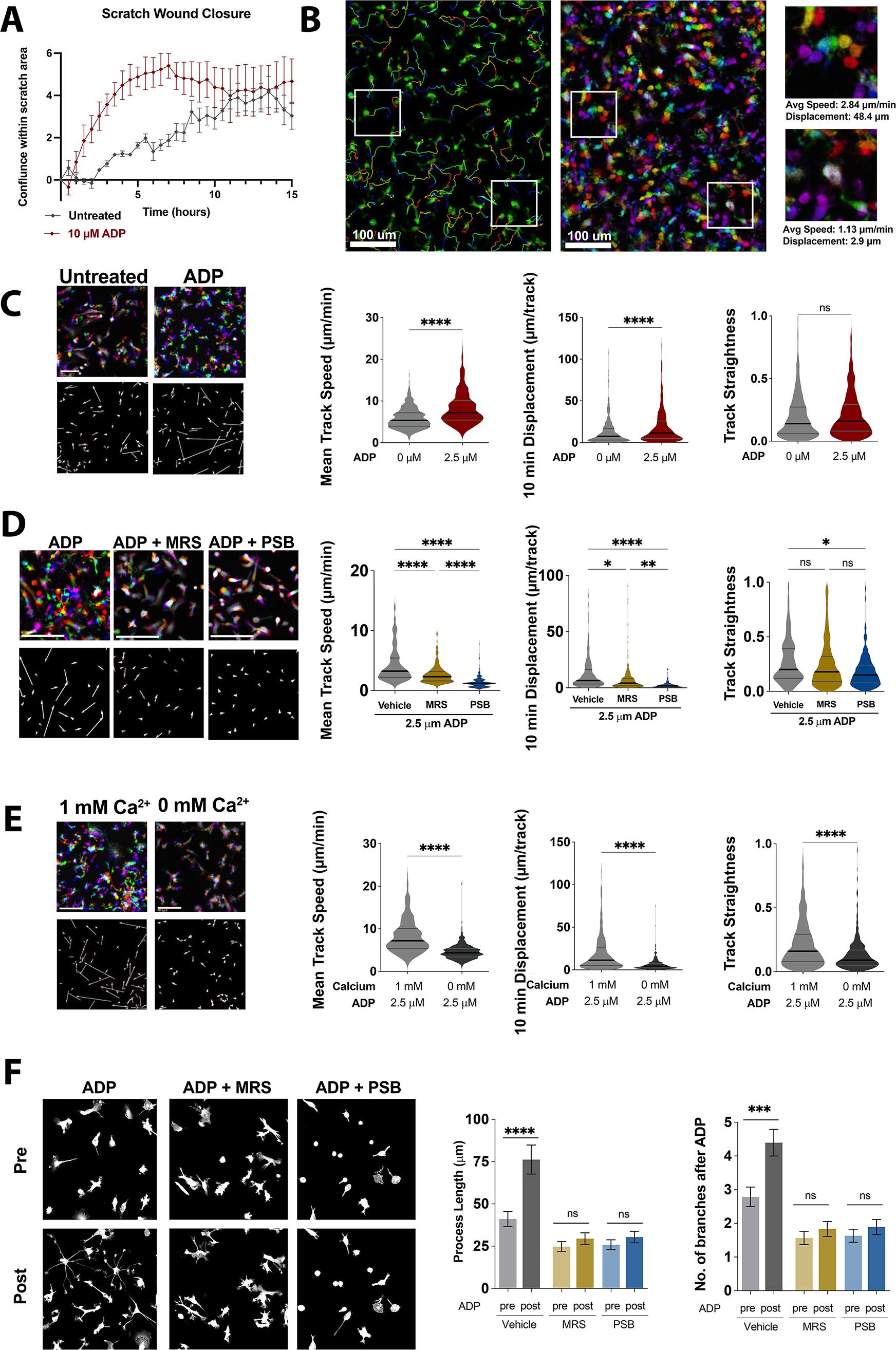
Nondirectional ADP exposure increases WT microglial speed and process extension. (**A**) Average trace showing closure of scratch wound produced with IncuCyte S3 WoundMaker. iPSC-microglia imaged every 30 min after scratch wound with or without ADP stimulation (n=4 wells; 2 images per well). (**B**) Representative image of WT iPSC-microglia motility 30 min after ADP exposure with cell tracks overlain (left). Pseudocolored images (center) across time: 0 min (red), 4 min (orange), 8 min (yellow), 12 min (green), 16 min (cyan), 20 min (blue), 24 min (purple), 28 min (magenta). Scale bar= 100 μm. White boxes zoomed in at right to demonstrate motile (top) and non-motile (bottom) cells. (**C**) Representative color images (top left) and displacement vectors (bottom left) of WT iPSC-microglia at baseline (no ADP, grey) and 30 min after 2.5 μM ADP treatment (red). Summary of Mean Speed (µm/min), Displacement over 10 min (μm/10 min) and Track straightness (track length/track displacement) (414-602 cells, 2 experiments). (**D**) Representative images, displacement vectors, and quantification of WT iPSC-microglia motility for 20 min following ADP addition. Cells were pre-treated with vehicle (grey), MRS 2211 (10 μM, gold), or PBS 0739 (10 μM, blue) (180-187 cells, 2 experiments). (**E**) Representative images, displacement vectors, and quantification of WT iPSC-microglia motility after ADP in 1 mM Ca^2+^ (light grey) or Ca^2+^-free buffer (dark grey) (401-602 cells, 3 experiments). (**F**) Representative images (left) and process extension (right) of iPSC-microglia (cytoplasmic GFP, grey) before or 30 min after ADP addition. Cells were pre-treated with vehicle (grey), MRS 2211 (10 μM, gold), or PBS 0739 (10 μM, blue) (52-163 cells, 3-4 experiments). (**C-F**) One way ANOVA with Tukey post hoc test. Data shown as mean ± SEM (A, F) and as violin plots with mean, 25^th^ and 75^th^ percentile (C-E). *P values* indicated by **ns** for non-significant, * for *P < 0.05, \**\* for *P < 0.01, \*\**\* for *P < 0.001* and **** for *P < 0.0001*.

In addition, some microglia responded to ADP by extending processes and altering their morphology rather than increasing motility (Figure 4-figure supplement 1). Microglia have been observed to extend processes in response to injury and purinergic stimulation in brain slices^8, 12^. Therefore, we compared process complexity before and 30 min after ADP exposure in WT microglia and observed significant increases in both the number of branches per process and total length of these processes (Figure 4F). Similar to effects on cell motility, ADP-mediated process extension was inhibited by P2Y_12_ and P2Y_13_ receptor antagonists (PSB 0739 and MRS 2211 respectively). Furthermore, even before process extension was activated with ADP, cells treated with P2Y antagonists showed significantly fewer and shorter processes, suggesting that baseline purinergic signaling may regulate resting microglial process dynamics. Altogether, these results demonstrate that activation of purinergic signaling through P2Y_12_ and P2Y_13_ receptors is required for ADP-driven microglial process extension and motility.

### ADP-evoked changes in cell motility and process extension are enhanced in TREM2-KO microglia

To characterize differences in motility characteristics between WT and TREM2 KO microglia responding to ADP, we plotted mean squared displacement (MSD) vs time and compared cell track overlays (flower plots) which showed that ADP enhances motility in TREM2 KO cells to a greater extent than in WT microglia (Figure 5A, B). Baseline motility characteristics in unstimulated cells, however, were similar in WT and TREM2 KO cells (Figure 5-figure supplement 1A, B). To further understand the basis of differences in ADP-induced motility between WT and TREM2 KO cells, we analyzed mean track speed, track displacement, and track straightness. Although mean track speeds were similar, TREM2 KO microglia showed greater displacement than WT cells (Figure 5C, D), raising the possibility that TREM2 KO cells may turn with lower frequency. Consistent with this, analysis of track straightness revealed that TREM2 KO microglia move farther from their origin for the same total distance traveled (Figure 5E). Vector autocorrelation, an analysis of directional persistence^62^, further confirmed that WT cells turn more frequently than TREM2 KO microglia in response to ADP (Figure 5-figure supplement 1C, D). To assess if these differences in TREM2 KO cells require sustained Ca^2+^ influx, we analyzed microglial motility in response to ADP stimulation in the absence of extracellular Ca^2+^ (Figure 5F-J). Mean-squared displacement (MSD) and cell-track overlay plots showed that motility is constrained when Ca^2+^ is removed from the external bath in both WT and TREM2 KO cells (Figure 5A, B **vs.** F, G). In the absence of extracellular Ca^2+^, TREM2 KO microglia showed similar mean speed, displacement, and track-straightness as WT cells (Figure 5C-E **vs** H-J). We conclude that increases in microglial motility (mean speed, displacement, and straightness) require sustained Ca^2+^ influx, and that deletion of TREM2 reduces microglial turning in response to ADP.

**Figure 5:**
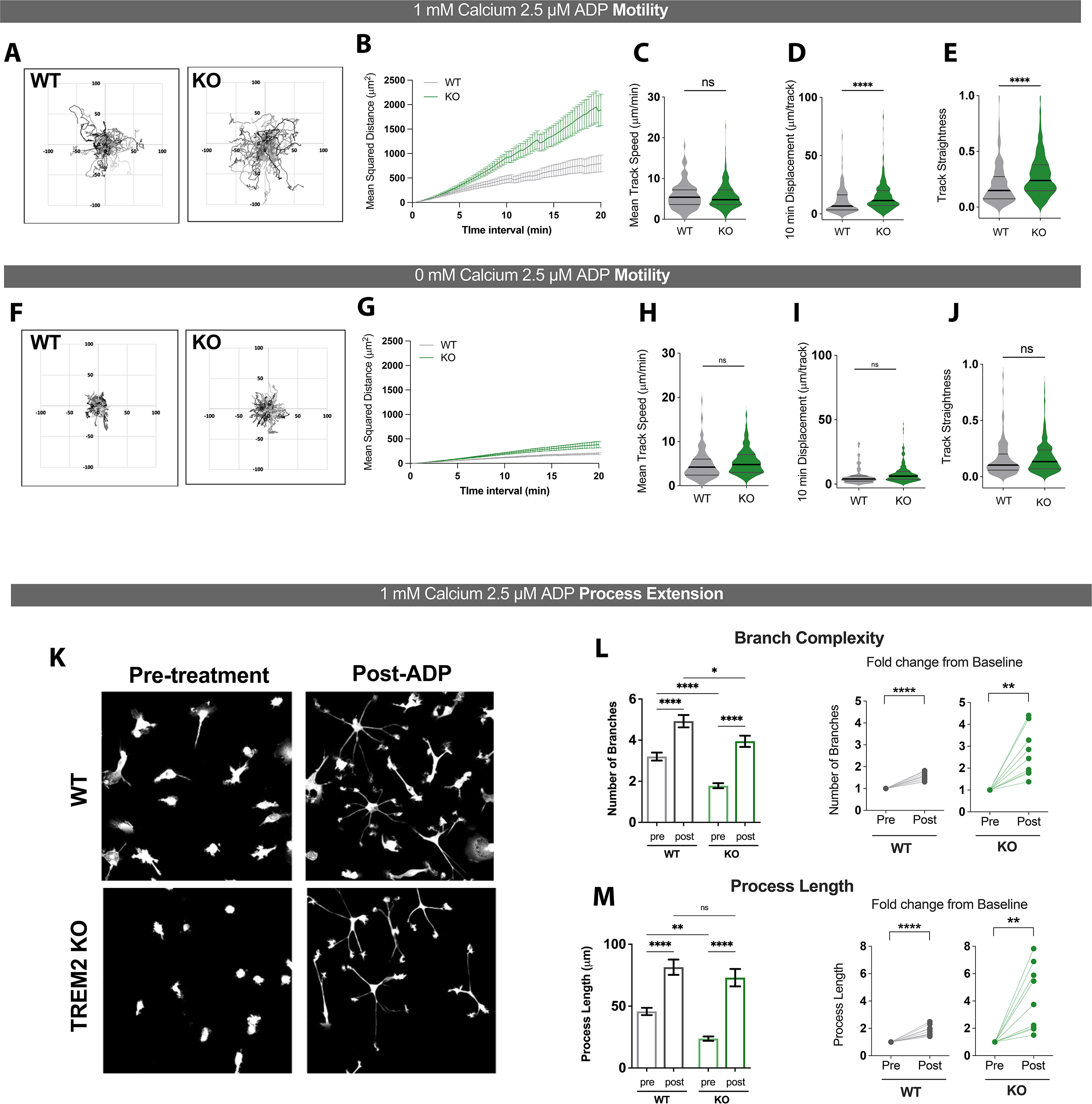
ADP-driven process extension and cell displacement are increased in TREM2 KO iPSC-microglia. (**A-E**) Motility of WT (grey) and TREM2 KO (green) iPSC-microglia over 20 min following ADP addition in 1 mM Ca^2+^-containing buffer. (**A**) Plots of track displacement in μm centered from point of origin at (0,0). (**B**) Mean-squared displacement (MSD) vs time. Mean-cell track speeds (**C**), total track displacement in 10-min interval (**D**), and track straightness (**E**) for 130-327 cells, 7 experiments, student’s t-test. (**F-J**). Same as (**A-F**) but in Ca^2+^-free medium (125-279 cells, 2 experiments, student’s t-test). (**K**) Representative images of GFP-expressing WT (top) and TREM2 KO (bottom) iPSC-microglia, before and 30 min after 2.5 μM ADP addition. (**L**) Quantification of total number of branches per cell before and after ADP treatment (left) and paired dot-plots showing fold change in branch number from pre-ADP levels (right). Each data-point represents an imaging field in the paired-plots. (**M**) Total process length before and after ADP treatment displayed as raw values per cell (left) and as fold change from baseline conditions per imaging field (right). For **L** and **M,** n=151-158 cells, WT; 133-167 cells, KO; 9-10 imaging fields, 3-4 experiments. One-way ANOVA with multiple comparisons for single-cell data, two-tailed paired t-test for the paired-plots. Data shown as mean ± SEM (B, G, L, M) and as violin plots with mean, 25^th^ and 75^th^ percentile (C-E, H-J). *P values* indicated by **ns** for non-significant, * for *P < 0.05, \**\* for *P < 0.01* and **** for *P < 0.0001*.

We next analyzed the effects of TREM2 deletion on process-extension in microglia. Treatment with ADP induced a dramatic increase in the number of branches and length of processes extended in both WT and TREM2 KO microglia (Figure 5K, L). Comparison of the absolute number of branches and process length after ADP treatment, as well as the relative fold-increase in these parameters from baseline indicated that process extension is not affected in TREM2 KO microglia (Figure 5K-M, figure 5-figure supplement 2A, B). We note that the greater fold-change in process extension in TREM2 KO cells can be attributed to the reduced morphological complexity of these cells prior to stimulation. Finally, ADP stimulation in Ca^2+^-free medium did not induce process extension in WT cells, and only a modest increase in TREM2 KO cells (Figure 5-figure supplement 2A **and** B **vs** C **and** D). Together, these results indicate that sustained Ca^2+^ entry across the PM is required for optimal microglial process extension in both WT and TREM2 KO microglia.

### Cytosolic Ca^2+^ levels tune motility in TREM2 KO iPSC-microglia

To further characterize the effects of sustained Ca^2+^ signals on microglial motility, we used Salsa6f-expressing iPSC WT and TREM2 KO reporter lines to monitor cytosolic Ca^2+^ and motility simultaneously in individual cells (Figure 6-figure supplement 1). To isolate the effects of sustained Ca^2+^ elevations on microglia motility, and eliminate any contribution from Ca^2+^ independent signaling pathways, we used a protocol that relies on triggering SOCE and varying external Ca^2+^ to maintain cytosolic Ca^2+^ at “low” or “high” levels in the Salsa6f reporter line (Figure 6A-C), similar to our previous study in T lymphocytes^63^. In WT cells, lowering extracellular Ca^2+^ from 2 to 0.2 mM predictably decreased the G/R ratio but did not influence mean track speed, 10-minute track displacement, or track straightness (Figure 6C, D **top**). However, in TREM2 KO microglia, reducing Ca^2+^ to a lower level significantly increased speed, displacement, and track straightness (Figure 6C**, D bottom**). These data suggest that motility characteristics of TREM2 KO microglia are more sensitive to changes in cytoplasmic Ca^2+^ levels than in WT cells. Similar results were obtained upon addition of ADP in this paradigm, suggesting that long-lasting Ca^2+^ elevations may override effects of Ca^2+^-independent ADP signaling on cell motility (Figure 6-figure supplement 2A).

**Figure 6:**
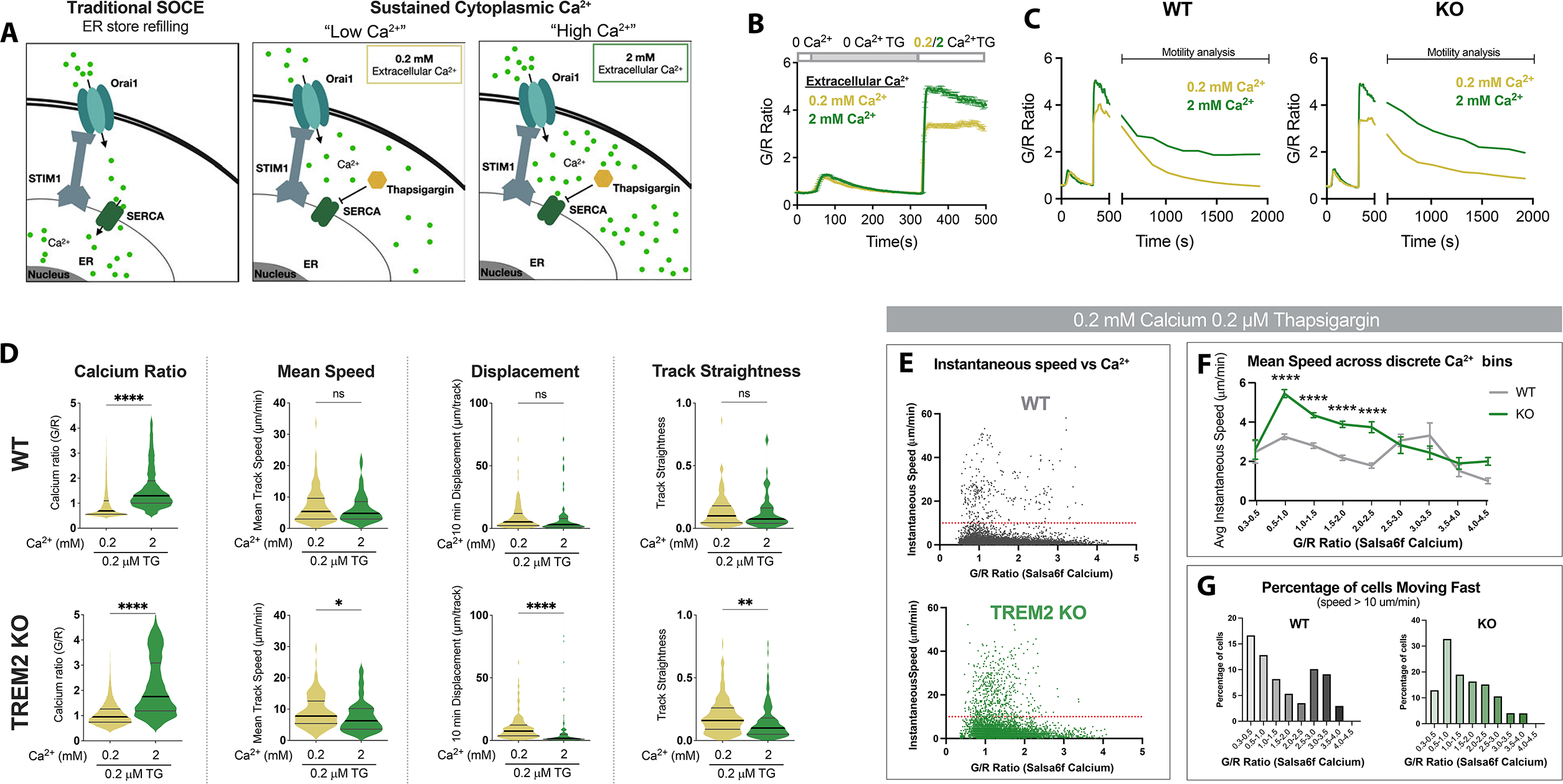
Cytosolic Ca^2+^ levels tune microglial motility in TREM2 KO cells. (**A**) Schematic of traditional SOCE pathway with store-refilling (left) and protocol for sustaining cytoplasmic Ca^2+^ to “low” and “high” levels with 0.2 and 2mM extracellular Ca^2+^ and using TG to inhibit store-refilling (right). (**B**) Average SOCE traces in WT Salsa6f iPSC-microglia showing changes in cytoplasmic Ca^2+^ after addition of either 0.2 or 2 mM extracellular Ca^2+^(n=78-110 cells). (**C**) Average change in cytoplasmic Ca^2+^ levels in WT and TREM2 KO microglia over 25 min after SOCE activation. (**D**) Comparison of Ca^2+^ levels and microglia motility in WT (top) and TREM2 KO (bottom) microglia. Cytosolic Ca^2+^ levels indicated by instantaneous single-cell G/R Ratio (n=74-158 cells). Mean of instantaneous speeds, track displacement and track straightness calculated as before in Figures 3 and 4. Yellow (0.2 mM Ca, TG), green (2 mM Ca, TG). Students t-test **** p < 0.0001; ** p = 0.0062; * p = 0.432; ns > 0.9999. (**E**) Correlation of instantaneous Ca^2+^ and instantaneous speed in WT and KO cells. Red line denotes 10 μm/sec (cells above this threshold considered “fast-moving”). For WT: p < 0.0001; r = −0.1316; number pairs = 5850. For KO: p < 0.0001; r = −0.1433; number pairs = 6063 (Spearman’s correlation). (**F**) Mean speed of cells binned by instantaneous G/R Ca^2+^ ratio (1-way ANOVA **** p < 0.0001). Each data point is calculated for a bin increment of 0.5 G/R ratio. (**G**) Percentage of fast-moving cells quantified as a function of G/R Ca^2+^ ratio. X-axis G/R ratios binned in increments of 0.5 as in **F**. In **E-G**, n=78-100 cells. Data shown as mean ± SEM (B, F) and as violin plots with mean, 25^th^ and 75^th^ percentile (D). *P values* indicated by **ns** for non-significant, * for *P < 0.05, \**\* for *P < 0.01* and **** for *P < 0.0001*.

To further analyze the Ca^2+^ dependence of microglial motility, we plotted Salsa6f G/R Ca^2+^ ratios for each individual cell at every time point against the instantaneous speeds of that cell (Figure 6E). These data revealed a stronger dependence of instantaneous speed on Ca^2+^ levels in TREM2 KO microglia (Figure 6F). Furthermore, when stratifying cell speed arbitrarily as “fast” (> 10 μm/min) or “slow” (< 10 μm/min), we observe a marked reduction in the percentage of “fast” cells when Ca^2+^ levels are high in TREM2 KO microglia (Figure 6G). Interestingly, frame-to-frame cell displacement correlated with cytosolic Ca^2+^ to the same degree in both WT and KO cells (Figure 6-figure supplement 2B, C). Together, TREM2 KO human microglia are more sensitive to tuning of motility by cytosolic Ca^2+^ than WT cells.

### Chemotactic defects in TREM2 KO microglia are rescued by dampening purinergic receptor activity

To assess the physiological significance of TREM2 deletion on microglial motility over longer time scales, we performed a scratch wound assay. At baseline, both WT and TREM2 KO microglia migrated into the cell free area at similar rates, consistent with our previous findings^41^ (Figure 7-figure supplement 1). Addition of ADP to this system accelerated the scratch wound closure rates to the same extent in WT and TREM2 KO. *In vivo,* directed migration of microglia is often driven by gradients of ADP from dying or injured cells^12, 30^. Because no chemical gradient is formed in the scratch wound assay^64^, we studied microglial chemotaxis toward ADP over a stable gradient using two-chamber microfluidic devices. Consistent with previous findings, WT iPSC-microglia directionally migrated up the concentration gradient of ADP resulting in higher numbers of cells within the central chamber^41, 65^. In the absence of a chemotactic cue, this directional migration was lost (Figure 7A). This assay revealed a deficit of chemotaxis in TREM2 KO microglia (Figure 7A), mirroring reports that TREM2 KO microglia are unable to migrate toward amyloid plaques in AD^40, 41, 66^. Given that ADP hypersensitivity in TREM2 KO cells is driven by increased expression of P2Y receptors, we examined the effects of dampening P2Y signaling to WT levels. Treatment with the P2Y_12_ receptor antagonist, PSB 0739, reduced Ca^2+^ responses in TREM2 KO cells and rescued the migration deficit in the chemotaxis assay (Figure 7B-C). These results link the increased Ca^2+^ signals and altered motility characteristics evoked by ADP in TREM2 KO cells to microglial chemotaxis toward areas of tissue damage, a vital functional response in microglia.

**Figure 7:**
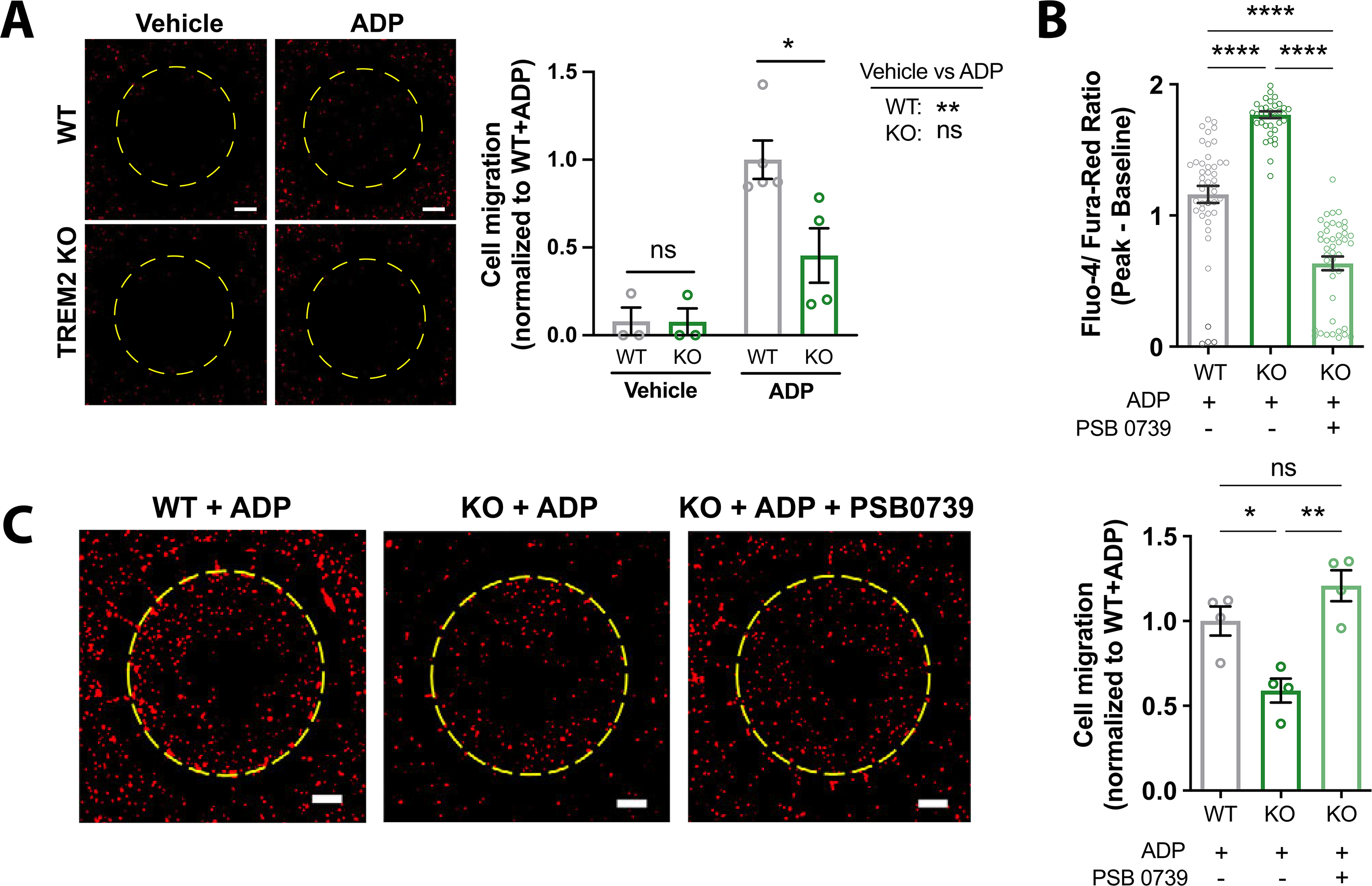
Migration deficits in TREM2 KO microglia are rescued by inhibition of purinergic signaling. (**A**) Migration towards ADP in a two-chamber microfluidic device. Representative images of RFP expressing microglia that migrated into the central chamber 3 days after 100 ng/mL ADP addition. Dotted circle delineates separation of inner and outer chamber. Scale bar = 500 μm. Quantification of microglial migration (right panel). Migrated cell counts are normalized to WT cells treated with ADP (n=3-4 experiments; One-way ANOVA with multiple comparisons). (**B**) Baseline subtracted peak ratiometric Ca^2+^ signal in response to 2.5 μM ADP in 1 mM extracellular Ca^2+^, and in the presence or absence of 10 μM PSB 0739 (44 cells, WT; 39-43 cells, KO; representative of 3 independent experiments; One-way ANOVA with multiple comparisons). (**C**) Two-chamber migration to 100 ng/mL ADP with or without 10 μM PSB 0739. Values are normalized to WT cells with ADP (n=3-4 experiments; One-way ANOVA with multiple comparisons). Representative images shown on the left. Scale bar = 500 μm. Data shown as mean ± SEM. *P values* indicated by **ns** for non-significant, * for *P < 0.05, \**\* for *P < 0.01* and **** for *P < 0.0001*.

## Discussion

This study focuses on two aims: understanding the roles of purinergic signaling in regulating human microglial motility behavior; and elucidating the impact of TREM2 loss of function on this Ca^2+^ signaling pathway. We find that sustained Ca^2+^ influx in response to ADP regulates microglial process extension, motility speed, and turning behavior. A key observation in our study is that microglia lacking TREM2 are highly sensitive to ADP-mediated signaling and show exaggerated cytoplasmic Ca^2+^ responses. Using novel iPSC-microglia lines that express a ratiometric, genetically encoded Ca^2+^ probe, Salsa6f, we found that the motility characteristics of human wild-type and TREM2-knockout microglia are differentially tuned by Ca^2+^ signaling. Informed by these discoveries, we were able to rescue chemotactic deficiencies in TREM2-knockout microglia by dampening purinergic receptor signaling.

We provide several lines of evidence to show that hyper-responsiveness to purinergic ADP signaling in TREM2 KO microglia is driven primarily by increased purinergic P2Y_12_ and P2Y_13_ receptor expression: (1) Ca^2+^ response is completely abrogated in the presence of P2Y_12_ and P2Y_13_ receptor inhibitors; (2) RNA-sequencing data shows significant increase in expression of P2Y_12_ and P2Y_13_ receptor transcripts but minimal fold-change in other regulators of Ca^2+^ signaling (IP3R, STIM, Orai, SERCA and PMCA); and (3) labelling of surface P2Y_12_ receptors shows greater PM expression in the TREM2 KOs. Furthermore, functional assays rule out any role for Ca^2+^ clearance mechanisms or any difference in maximal IP_3_ and SOCE activity as a cause of increased sustained Ca^2+^ signal in TREM2 KO cells. Mechanistically, this increase in Ca^2+^ signals is driven by enhanced IP_3_-mediated ER store-release coupled to SOCE. Indeed, based on the dose-response curves for peak ADP-Ca^2+^ responses in Ca^2+^ free buffer, TREM2 KO cells have an EC_50_ at least 10-fold lower than WT cells. As a functional consequence, TREM2 KO microglia exhibit a defect in turning behavior, and show greater displacement over time despite moving with similar speeds as the WT cells. The increased frequency in turning in WT microglia (relative to TREM KO cells) reflects greater canceling of the velocity vectors, which take the direction of motility into account. This restricts cell motility to more confined regions, potentially allowing for more frequent path correction. It is important to note that these motility differences with ADP are observed after acute treatment and in the absence of any gradient.

Interestingly, deletion of TREM2 had no significant impact on scratch wound closure rates, over a time scale of 24 hours in the presence of a constant concentration of ADP^67^. However, we find in a directional chemotaxis assay towards a gradient of ADP concentration that TREM2 KO cells are unable to migrate as efficiently as WT cells, concordant with previous studies showing reduced migration of TREM2 KO cells towards Aβ plaques^41^. Enhanced ADP signaling likely abolishes the ability of TREM2 KO cells to distinguish gradations of the agonist, and this loss of gradient-sensing results in an inability to perform directed migration. We speculate that increased ADP Ca^2+^ signaling in TREM2 KO cells may result in Ca^2+^ signaling domains that are no longer restricted to the cell region near to the highest ADP concentrations, and disrupt the polarity of key signaling molecules that drive directed cell motility.

The amplitude and duration of Ca^2+^ signals shape specificity of downstream cellular responses. Our experiments with ADP in Ca^2+^ free medium revealed that a transient Ca^2+^ signal is insufficient to induce microglial motility in either WT or TREM2 KO cells. Previous studies have shown that mouse microglia with genetic deletion of STIM1 or Orai1 also show defects in cell migration to ATP^57, 68^, likely because diminished SOCE renders them unable to sustain Ca^2+^ signals in response to ATP. The dependence of motility on prolonged purinergic Ca^2+^ signals may thus be a general feature of microglia. In contrast, a Ca^2+^ transient can initiate some process extension in TREM2 KO but not in WT microglia, suggesting a threshold for ADP signaling that is reached in KO but not WT cells, and highlighting subtle differences in the Ca^2+^ requirement for motility and process extension in TREM2 KO microglia.

To directly monitor Ca^2+^ signaling and motility simultaneously in individual cells, we developed a novel iPSC-microglia cell-line expressing a genetically encoded, ratiometric Ca^2+^ indicator Salsa6f, a GCaMP6f-tdTomato fusion protein. Because Salsa6f allows simultaneous measurement of Ca^2+^ signal and tracking of processes, this Salsa6f iPSC line is likely to be a useful tool to dissect the relationship between Ca^2+^ signaling and the function of various iPSC-derived human cell types including neurons, astrocytes, and microglia. In addition, this line may be readily xenotransplanted for use with human/microglia chimeric models to examine functional Ca^2+^ responses to injury and pathology *in vivo*. Using Salsa6f-expressing microglia, we uncovered critical differences in how Ca^2+^ levels tune motility in WT and TREM2 KO microglia. By tracking instantaneous velocity at the same time as Salsa6f Ca^2+^ ratios in individual cells, we found that TREM2 KO cell motility showed a greater sensitivity to changes in cytosolic Ca^2+^ levels with significantly higher speeds than WT cells at lower Ca^2+^ and a more dramatic reduction in cell speed at high Ca^2+^ levels. It is possible that high cytosolic Ca^2+^ serves as a temporary STOP signal in microglia similar to its effects on T cells ^63^; we further speculate that TREM2 KO cells may be more subject to this effect with ADP, given the higher expression of P2RY_12_ and P2Y_13_ receptors. Accordingly, reducing cytosolic Ca^2+^, resulted in increased mean speed, displacement, and straighter paths for TREM2 KO iPSC-microglia, but had no effect on these motility metrics in WT cells suggesting that TREM2 KO cells may display a greater dynamic range in regulating their motility in response to sustained Ca^2+^ elevations. Consistent with this observation, chemotaxis in TREM2 KO cells was restored by partially inhibiting P2Y_12_ receptors. In response to neurodegenerative disease, microglia down-regulate P2Y_12_ receptors^23, 25, 31^. Active regulation of purinergic receptor expression is critical for sensing ADP gradients and decreasing motility near the chemotactic source. *In vivo* studies^21, 23, 41^ suggest that TREM2 KO microglia are unable to down-regulate P2Y receptor expression upon activation, which may lead to the known chemotactic deficits in these cells.

The studies presented here provide evidence that reducing purinergic receptor activity may be clinically applicable in Alzheimer’s patients with TREM2 loss of function mutations ^40, 55, 69^. Pharmacologically targeting P2Y_12_ receptors to dampen both the Ca^2+^ dependent (PLC) and independent (DAG) arms of the GPCR signaling pathway may be useful to control microglial activation and motility. However, our results suggest that altering downstream Ca^2+^ flux may be sufficient, and thus, CRAC (Orai1) channel blockers that would specifically inhibit the sustained Ca^2+^ signals without affecting the initial Ca^2+^ transient or the activation of DAG may provide a more targeted approach.

Currently, TREM2 activating antibodies are being examined in early stage clinical trials for Alzheimer’s disease^70, 71^, making it critically important to understand the broad consequences of TREM2 signaling. Therefore, an understanding of how TREM2 influences responses to purinergic signals and regulates cytosolic Ca^2+^ in human iPSC-microglia is critical. Beyond TREM2, we have found that protective variants in MS4A6A and PLCG2 gene expression also decrease P2Y_12_ and P2Y_13_ receptor expression (unpublished data), suggesting this mechanism of microglial activation could be common across several microglial AD risk loci.

In summary, deletion of TREM2 renders iPSC-microglia highly sensitive to ADP, leading to prolonged Ca^2+^ influx which increases cell displacement by decreasing cell turning. Despite this, TREM2 KO microglia show a defect in chemotaxis that is likely due to their inability to sense ADP gradients and make appropriate course corrections. Decreasing purinergic signaling in TREM2 KO microglia rescues directional chemotactic migration. We suggest that purinergic modulation or direct modulation of Ca^2+^ signaling could provide novel therapeutic strategies in many AD patient populations, not solely those with reduced TREM2 function.

## Materials and Methods

### Key Resources Table

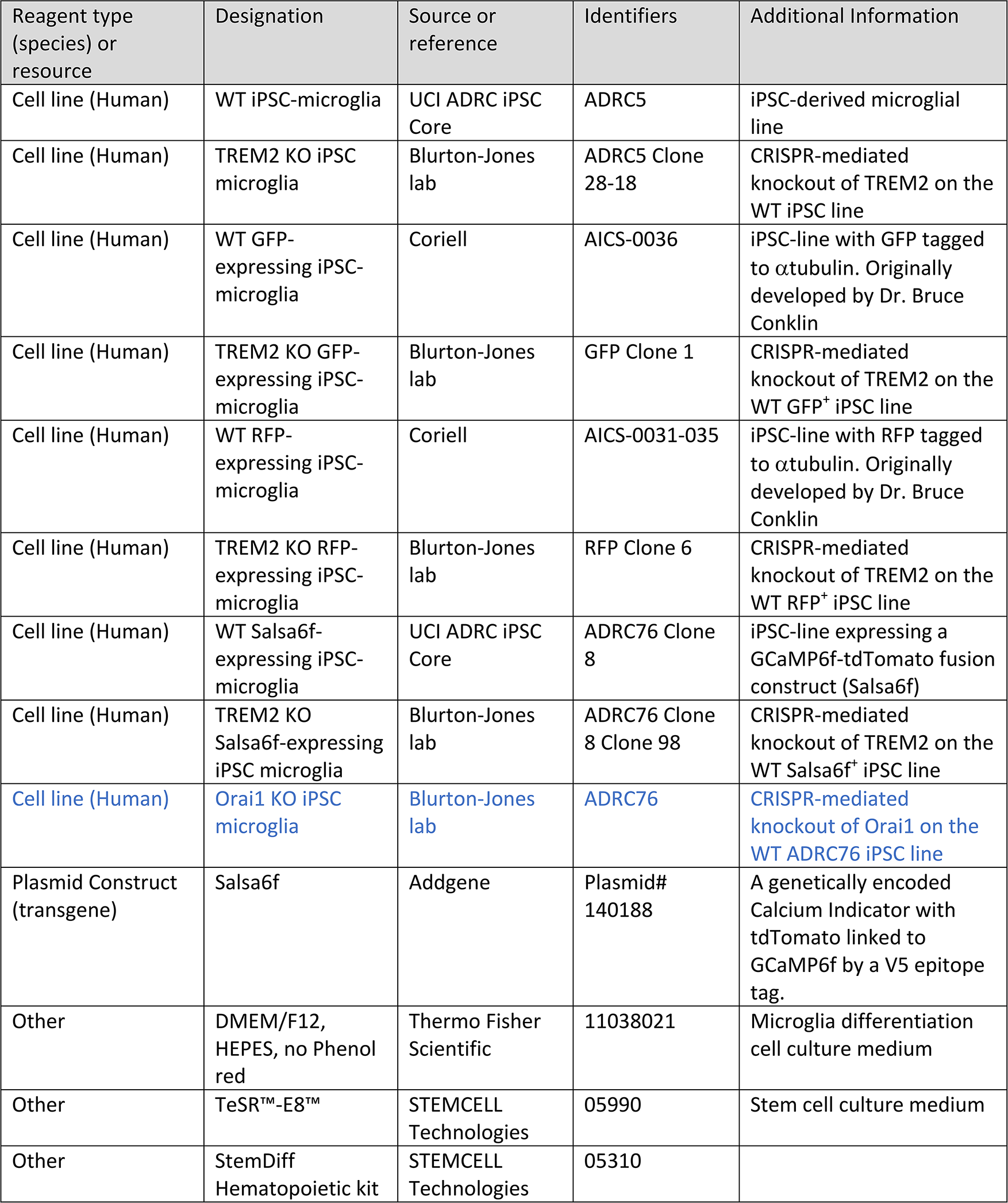
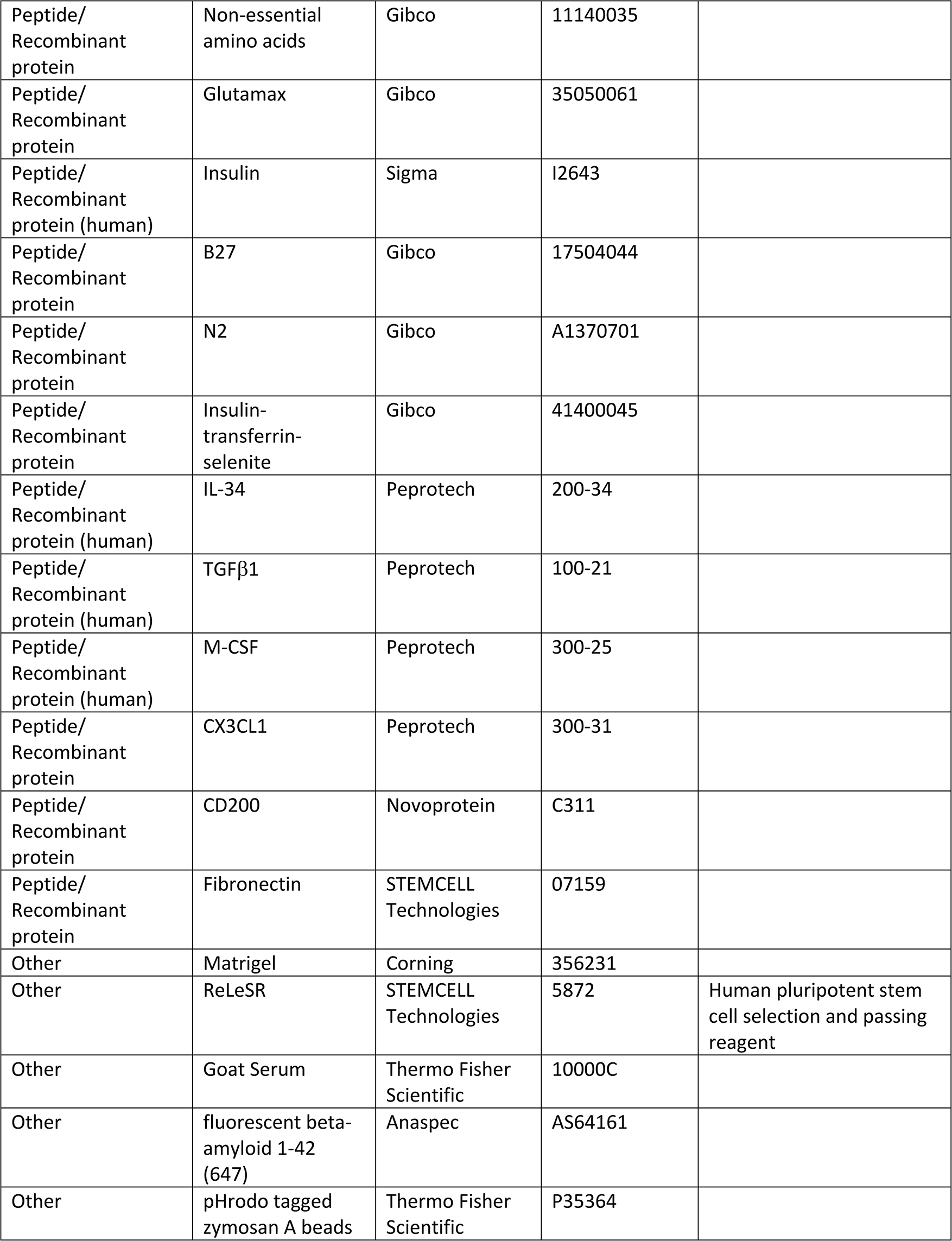
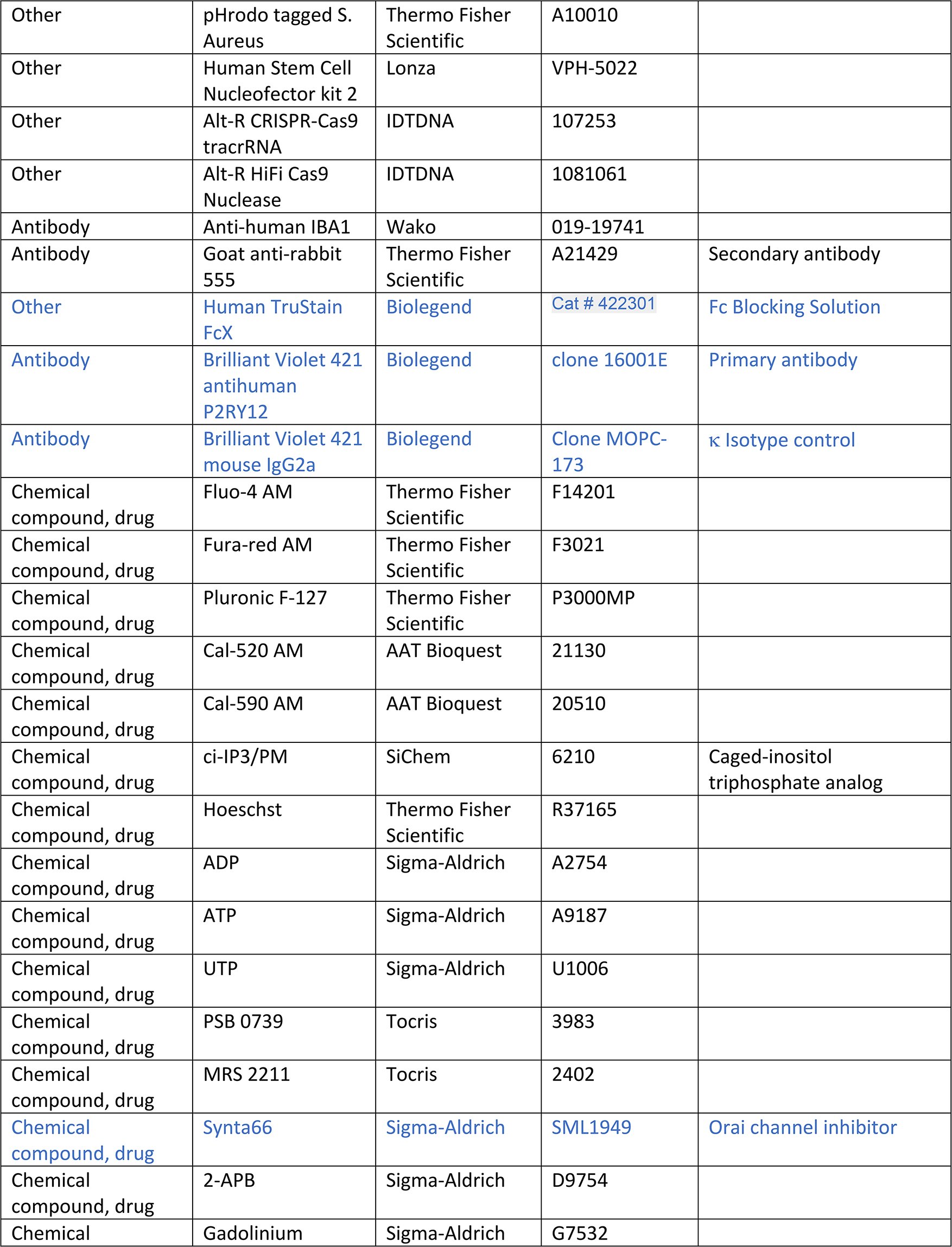
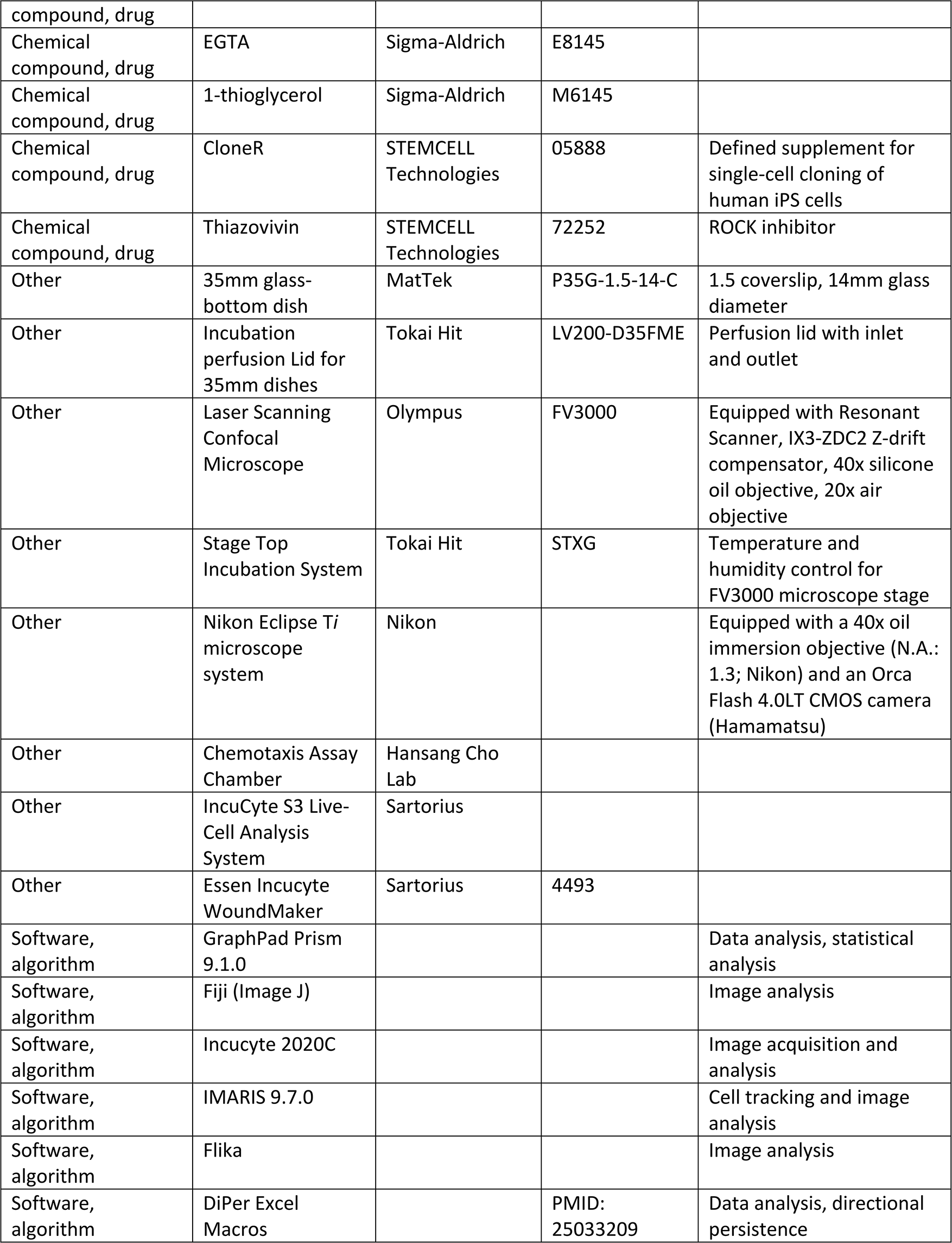
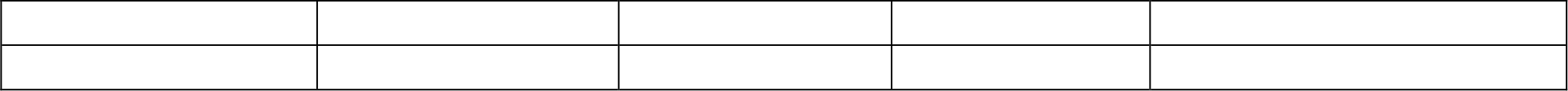

#### Generation of iPSCs from human fibroblasts

Human induced pluripotent stem cell lines were generated by the University of California, Irvine Alzheimer’s Disease Research Center (UCI ADRC) Induced Pluripotent Stem Cell Core from subject fibroblasts under approved Institutional Review Boards (IRB) and human Stem Cell Research Oversight (hSCRO) committee protocols. Informed consent was received from all participants who donated fibroblasts. Reprogramming was performed with non-integrating sendai virus in order to avoid integration effects. To validate new iPSC lines, cells were karyotyped by G-banding and tested for sterility. Pluripotency was verified by Pluritest Array Analysis and trilineage in vitro differentiation. Additional GFP- and RFP-αtubulin expressing iPSC lines (AICS-0036 and AICS-0031-035) were purchased from Coriell and originally generated by Dr. Bruce Conklin. iPSCs were grown antibiotic free on Matrigel (Corning) in complete mTeSR1 or TeSR-E8 medium (STEMCELL Technologies) in a humidified incubator (5% CO_2_, 37° C). All lines will be available upon request to the corresponding author.

#### CRISPR-mediated knockout of TREM2 and ORAI1

Genome editing to delete TREM2 was performed as in McQuade et al. 2020^41^. Briefly, iPSCs were nucleofected with Ribonucleoprotien complex targeting the second exon of TREM2 and allowed to recover overnight. Transfected cells were dissociated with pre-warmed Accutase then mechanically plated to 96-well plates for clonal expansion. Genomic DNA from each colony was amplified and sequenced at the cut site. The amplification from promising clones was transformed via TOPO cloning for allelic sequencing. Knockout of TREM2 was validated by western blotting (AF1828, R&D) and HTRF (Cisbio) ^41^. A similar strategy was used to delete ORAI1 using an RNP complex of Cas9 protein coupled with a guide RNA (5’ CGCTGACCACGACTACCCAC) targeting the second exon of ORAI1. All iPSC lines were confirmed to be sterile and exhibiting a normal Karyotype via Microarray-based Comparative Genomic Hybridization (aCGH, Cell Line Genetics).

#### iPSC-microglia differentiation

iPSC-microglia were generated as described in ^50, 51^. Briefly, iPSCs were directed down a hematopoetic lineage using the STEMdiff Hematopoesis kit (STEMCELL Technologies). After 10-12 days in culture, CD43+ hematopoteic progenitor cells are transferred into a microglia differentiation medium containing DMEM/F12, 2× insulin-transferrin-selenite, 2× B27, 0.5× N2, 1× Glutamax, 1× non-essential amino acids, 400 μM monothioglycerol, and 5 μg/mL human insulin. Media was added to cultures every other day and supplemented with 100 ng/mL IL-34, 50 ng/mL TGF-β1, and 25 ng/mL M-CSF (Peprotech) for 28 days. In the final 3 days of differentiation 100 ng/mL CD200 (Novoprotein) and 100 ng/mL CX3CL1 (Peprotech) were added to culture.

#### Confocal Laser Scanning Microscopy

Unless otherwise stated, cells were imaged on an Olympus FV3000 confocal laser scanning inverted microscope equipped with high-speed resonance scanner, IX3-ZDC2 Z-drift compensator, 40x silicone oil objective (NA 1.25) and a Tokai-HIT stage top incubation chamber (STXG) to maintain cells at 37°C. To visualize Salsa6f, 488 nm and 561 nm diode lasers were used for sequential excitation of GCaMP6f (0.3% laser power, 450V channel voltage, 494-544nm detector width) and TdTomato (0.05% laser power, 450V channel voltage, 580-680nm detector width), respectively. Fluo-4 and Fura-red were both excited using a 488 nm diode laser (0.07% laser power, 500V channel voltage, 494-544nm detector width for Fluo-4; 0.07% laser power, 550V channel voltage, 580-680nm detector for Fura-Red). Two high-sensitivity cooled GaAsP PMTs were used for detection in the green and red channels respectively. GFP was excited using the same settings as GCaMP6f. Other image acquisition parameters unique to Ca^2+^ imaging, microglia process and cell motility analysis are indicated in the respective sections.

### Measurement of intracellular Ca^2+^

#### Cell preparation

iPSC-microglia were plated on fibronectin-coated (5 μg/mL) glass-bottom 35 mm dishes (MatTek, P35G-1.5-14-C) overnight at 60 % confluence. Ratiometric Ca^2+^ imaging was done using Fluo-4 AM and Fura-Red AM dyes as described previously^41^. Briefly, cells were loaded in microglia differentiation medium with 3 μM Fluo-4 AM and 3 μM Fura-Red AM (Molecular Probes) in the presence of Pluronic Acid F-127 (Molecular Probes) for 30 min at room temperature (RT). Cells were washed with medium to remove excess dye and 1 mM Ca^2+^ Ringer’s solution was added to the 35 mm dish before being mounted on the microscope for live cell imaging. We note that iPSC-microglia are sensitive to shear forces and produce brief Ca^2+^ signals in response to solution exchange that are dependent on extracellular Ca^2+^, and that these are more prominent at 37° C. To minimize these confounding effects, cells were imaged at RT and perfusion was performed gently. Salsa6f-expressing iPSC-microglia were prepared for Ca^2+^ imaging in the same way as conventional microglia, but without the dye loading steps. The following buffers were used for Ca^2+^ imaging: (1) 1 or 2 mM Ca^2+^ Ringer solution comprising 155 mM NaCl, 4.5 mM KCl, 1 mM CaCl_2_, 0.5 mM MgCl_2_, 10 mM glucose, and 10 mM HEPES (pH adjusted to 7.4 with NaOH), (2) Ca^2+^-free Ringer solution containing: 155 mM NaCl, 4.5 mM KCl, 1.5 mM MgCl_2_, 10 mM glucose, 1 mM EGTA, 10 mM HEPES, pH 7.4. Live cell imaging was performed as described earlier. Cells were treated with ADP as indicated in the results section. *Data acquisition:* Time-lapse images were acquired in a single Z-plane at 512 x 512 pixels (X = 318.2 μm and Y = 318.2 μm) and at 2-3 sec time intervals using Olympus FV3000 software. Images were time averaged over 3 frames to generate a rolling average and saved as .OIR files. *Data analysis:* Time-lapse videos were exported to Fiji-ImageJ (https://imagej.net/Fiji), converted to tiff files (16-bit) and background subtracted. Single-cell analysis was performed by drawing ROIs around individual cells in the field and average pixel intensities in the green and red channels were calculated for each ROI at each time-point. GCaMP6f/ TdTomato (G/R Ratio) and Fluo-4/Fura-Red ratio was then obtained to further generate traces showing single-cell and average changes in cytosolic Ca^2+^ over time. Single-cell ratio values was used to calculate Peak Ca^2+^ signal and responses at specific time points after agonist application as previously reported^60^. Peak Ca^2+^ signal for each cell was baseline subtracted, which was calculated as an average of 10 minimum ratio values before application of agonist. SOCE rate was calculated as Δ(Ratio)/ Δt (sec^-1^) over a 10-sec time frame of maximum initial rise after Ca^2+^ add-back. Area under the curve (AUC) was calculated using the AUC function in GraphPad Prism.

### Microglia process extension analysis

#### Data acquisition

GFP-expressing iPSC-microglia were plated overnight on 35 mm glass bottom dishes at 40-50% confluence. Cells were imaged by excitation of GFP on the confocal microscope at 37° C as described earlier. To study process extension in response to ADP, two sets of GFP images were obtained for each field of view across multiple dishes: before addition of ADP (baseline) and 30 min after application of ADP. Images were acquired as a Z-stack using the Galvo scanner at Nyquist sampling. Adjacent fields of view were combined using the Stitching function of the Olympus FV3000 Software and saved as .OIR files.

#### Process Analysis

The basic workflow for microglia process analysis was adapted from Morrison et al, Sci. Rep, 2017 ^72^. Image stacks (.OIR files) were exported to Fiji-Image J and converted into 16-bit Tiff files using the Olympus Viewer Plugin (https://imagej.net/OlympusImageJPlugin). Maximum intensity projection (MIP) image from each Z-stack was used for further processing and analysis. MIP images were converted to 8-bit grey scale images, to which a threshold was applied to obtain 8-bit binary images. The same threshold was used for all sets of images, both before and after ADP application. Noise reduction was performed on the binary images using the Process -> Noise -> Unspeckle function. Outlier pixels were eliminated using Process -> Noise -> Outliers function. The binary images were then skeletonized using the Skeletonize2D/3D Plugin for Image J (https://imagej.net/plugins/skeletonize3d). Sparingly, manual segmentation was used to separate a single skeleton that was part of two cells touching each other. The Analyze Skeleton Plugin (https://imagej.net/plugins/analyze-skeleton/) was then applied to the skeletonized images to obtain parameters related to process length and number of branches for each cell in the imaging field. Processes were considered to be skeletons > 8 μm. The data was summarized as average process length and number of branches, before and after ADP application for a specific imaging field, normalized to the number of cells in the field which allowed for pairwise comparison. Additionally, single cell data across all experiments were also compared in some instances.

#### IP_3_ uncaging

Whole-field uncaging of i-IP_3_, a poorly metabolized IP_3_ analog, was performed as previously described ^73^ with minor modifications. Briefly, iPSC-microglia were loaded for 20 min at 37° C with either Cal520 AM or Cal590 AM (5 μM, AAT Bioquest), and the cell permeable, caged i-IP_3_ analog ci-IP_3_/PM (1 μM, SiChem) plus 0.1% Pluronic F-127 in Microglia Basal Medium. Cells were washed and incubated in the dark for further 30 min in a HEPES-buffered salt solution (HBSS) whose composition was (in mM): 135 NaCl, 5.4 KCl, 1.0 MgCl2, 10 HEPES, 10 glucose, 2.0 CaCl_2_, and pH 7.4. Intracellular Ca^2+^ ([Ca^2+^]_i_) changes were imaged by employing a Nikon Eclipse T*i* microscope system (Nikon) equipped with a 40x oil immersion objective (N.A.: 1.3; Nikon) and an Orca Flash 4.0LT CMOS camera (Hamamatsu). Cal520 or Cal590 were excited by a 488 or a 560 nm laser light source (Vortran Laser Technologies), respectively. i-IP3 uncaging was achieved by uniformly exposing the imaged cells to a single flash of ultraviolet (UV) light (350-400 nm) from a Xenon arc lamp. UV flash duration, and thus the amount of released i-IP_3_, was set by an electronically controlled shutter.

Image acquisition was performed by using Nikon NIS (Nikon) software. After conversion to stack tiff files, image sequences were analyzed with Flika, a custom-written Python-based imaging analysis software (https://flika-org.github.io/; ^74^). After background subtraction, either Cal520 or Cal590 fluorescence changes of each cell were expressed as ΔF/F_0_, where F_0_ is the basal fluorescence intensity and ΔF the relative fluorescence change (F_x_ – F_0_). Data are reported as superplots ^75^ of at least three independent replicates. Experiments were reproduced with two independent lines. Comparisons were performed by unpaired non-parametric t-test.

#### Immunocytochemistry

Cells were fixed with 4 % paraformaldehyde for 7 min and washed 3x with 1X PBS. Blocking was performed at room temp for 1 hr in 5 % Goat Serum, 0.1 % Triton5 X-100. Primary antibodies were added at 1:200 overnight 4° C (IBA1, 019-19741, FUJIFILM Wako). Plates were washed 3x before addition of secondary antibodies (Goat anti-Rabbit 555, ThermoFisher Scientific) and Hoechst (ThermoFisher Scientific). Images were captured on an Olympus FV3000RS confocal microscope with identical laser and detection settings. Images were analyzed with IMARIS 9.7.0 software.

#### Flow Cytometry

iPSC-derived microglia were seeded on fibronectin-coated 12-well plates at 200,000 cells/well. Cells were harvested and centrifuged in FACS tubes at 300 xG for 5 min at 4° C. The cell pellet was subsequently resuspended in FACS buffer (1X PBS + 0.5% FBS). Fc receptors were blocked with a blocking buffer (Bio-legend TruStain FcX in 1X PBS + 10% FCS). Cells were then incubated with Brilliant Violet 421-labelled anti-human P2Y_12_ receptor antibody (clone S16001E, Biolegend, Cat# 392106) or with IgG2a isotype control antibody (clone MOPC-173, Biolegend, Cat# 400260) for 30 min at 4° C. Cells were washed, pelleted, and then resuspended in FACS buffer. Clone S16001E binds to the extracellular domain of the P2Y_12_ and permits labeling of plasma membrane P2Y_12_ receptors. Data were acquired using Novocyte Quanteon flow cytometer (Agilent) and analyzed using FlowJo analysis software (FlowJo v10.8.1 LLC Ashland, Oregon).

#### Scratch wound assay

Nondirectional motility was analyzed using Essen Incucyte WoundMaker. iPSC-microglia were plated on fibronectin (STEMCELL Technologies) at 90% confluence. Scratches were repeated 4x to remove all cells from the wound area. Scratch wound confluency was imaged every hour until scratch wound was closed (15 hrs). Confluence of cells within the original wound ROI was calculated using IncuCyte 2020C software.

#### IMARIS Cell Tracking

For motility assays, iPSC-microglia were tracked using a combination of manual and automatic tracking in IMARIS 9.7.0 software. For videos of GFP lines, cells were tracked using spot identification. For videos of Salsa6f lines, surface tracking was used to determine ratiometric Ca^2+^ fluorescence and motility per cell. In both conditions, tracks were defined by Brownian motion with the maximum distance jump of 4 microns and 10 frame disturbance with no gap filling. Tracks shorter than 3 minutes in length were eliminated from analysis. After automated track formation, tracks underwent manual quality control to eliminate extraneous tracks, merge falsely distinct tracks, and add missed tracks. After export, data was plotted in Prism 9.1.0 or analyzed in excel using DiPer macros for Plot_At_Origin (translation of each trajectory to the origin) and mean squared distance (MSD) MSD(t)=4D(t-P(1-e^(-t/P))) where D is the diffusion coefficient, t is time, and P represents directional persistence time (time to cross from persistent directionality to random walk)^62^. From IMARIS, speed was calculated as instantaneous speed of the object (mm/s) as the scalar equivalent to object velocity. These values were transformed to <u>m</u>m /min as this time scale is more relevant for the changes we observed. Mean track speed represents the mean of all instantaneous speeds over the total time of tracking. 10 min displacement is calculated by (600) * (TDL/TD), where TDL = track displacement length (distance between the first and last cell position) represented as TDL= p(n) - p(1) for all axes where the vector p is the distance between the first and last object position along the selected axis and TD = track duration represented as TD = T(n) - T(1), where T is the timepoint of the first and final timepoint within the track. Frame-to-frame displacement is calculated as p(n) – p(n-1) for all the different frames in a cell track. Track straightness is defined as TDL/TL where TDL = track displacement as described above and TL = track length representing the total length of displacements within the track TL= sum from t=2 to n of |p(t)-p(t-1)|.

#### Generation of Salsa6f-expressing iPSC lines

iPSCs were collected following Accutase enzymatic digestion for 3 min at 37° C. 20,000 cells were resuspended in 100 μL nucleofection buffer from Human Stem Cell Nucleofector™ Kit 2 (Lonza). Salsa6f-AAVS1 SHL plasmid Template (2 μg; Vector Builder) and RNP complex formed by incubating Alt-R® S.p. HiFi Cas9 Nuclease V3 (50 μg; IDTDNA) was fused with crRNA:tracrRNA (IDTDNA) duplex for 15 min at 23° C. This complex was combined with the cellular suspension and nucleofected using the Amaxa Nucleofector program B-016. To recover, cells were plated in TeSR™-E8™ (STEMCELL Technologies) media with 0.25 μM Thiazovivin (STEMCELL Technologies) and CloneR™ (STEMCELL Technologies) overnight. The following day, cells were mechanically replated to 96-well plates in TeSR™-E8™ media with 0.25 μM Thiazovivin and CloneR™ supplement for clonal isolation and expansion. Plates were screened visually with a fluorescence microscope to identify TdTomato^+^ clones. Genomic DNA was extracted from positive clones using Extracta DNA prep for PCR (Quantabio) and amplified using Taq PCR Master Mix (Thermo Fisher Scientific) to confirm diallelic integration of the Salsa6f cassette. A clone confirmed with diallelic Salsa6f integration in the AAVS1 SHL was then retargeted as previously described^41^ to knock-out Trem2.

#### Phagocytosis assay

Phagocytosis of transgenic iPSC-microglia was validated using IncuCyte S3 Live-Cell Analysis System (Sartorius) as in McQuade et al. 2020^41^. Microglia were plated at 50% confluency 24 hours before substrates were added. Cells were treated with 50 μg/mL pHrodo tagged human AD synaptosomes (isolated as described in McQuade et al. 2020), 100 ng/mL pHrodo tagged zymosan A beads (Thermo Fisher Scientific), 100 ng/mL pHrodo tagged S. Aureus (Thermo Fisher Scientific), or 2 μg/mL fluorescent beta-amyloid (Anaspec). Image masks for fluorescence area and phase were generated using IncuCyte 2020C software.

#### Chemotaxis assay

iPSC-microglia were loaded into the angular chamber (2-5K cells/device) to test activation and chemotaxis towards the central chamber containing either ADP (100 ng/mL or 234 nM) or vehicle. When noted, PSB 0739 (10 μM) was added to both the central and angular chamber to inhibit P2Y_12_ receptors. To characterize motility, we monitored the number of recruited microglia in the central chamber for 4 days under the fully automated Nikon TiE microscope (10× magnification; Micro Device Instruments, Avon, MA, USA).

### Statistical Analysis

GraphPad Prism (Version 6.01 and 8.2.0) was used to perform statistical tests and generate P values. We used standard designation of P values throughout the Figures (**ns**, not significant or *P ≥ 0.05*; * *P < 0.05*; ** *P < 0.01*; *** *P < 0.001*; **** *P < 0.0001*). Traces depicting average changes in cytosolic Ca^2+^ over time are shown as mean ± SEM (Standard Error of Mean). Accompanying bar-graphs with bars depicting mean ± SEM (Standard Error of Mean) provide a summary of relevant parameters (Amplitude of Ca^2+^ response, degree of store-release, rate of Ca^2+^ influx etc) as indicated. Details of number of replicates and the specific statistical test used are provided in the individual figure legends.

## Acknowledgements

The authors would like to thank Dr. Andy Yeromin for the development of Excel macros to analyze IMARIS cell tracking. The authors would also like to thank Morgan Coburn for sharing python scripts that aided in the organization of IMARIS output files. This work was supported by T32 NS082174 and ARCS foundation (A.M.); the European Union’s Horizon 2020 research and innovation program under the Marie Sklodowska-Curie grant agreement iMIND – No. 84166 (A.G.); NIH R01 NS14609 and AI121945 (M.D.C.); NIH U01 AI160397 (S.O.); NRF 2020R1A2C2010285, 2020M3C7A1023941, and NIH AG059236-01A1 (H.C.); NIH AG048099, AG056303, and AG055524 (M.B.J.); RF1DA048813 (M.B.J. and S.G.); UCI Sue & Bill Gross Stem Cell Research Center Seed Grant (S.G.); and a generous gift from the Susan Scott Foundation (M.B.J.). iPSC lines were generated by the UCI-ADRC iPS cell core funded by NIH AG066519. Experiments using the GFP-expressing iPSC line AICS-0036 were made possible through the Allen Cell Collection, available from Coriell Institute for Medical Research.

## Ethics

Human iPSC lines were generated by the University of California Alzheimer’s Disease Research Center (UCI ADRC) stem cell core. Subject fibroblasts were collected under approved Institutional Review Boards (IRB) and human Stem Cell Research Oversight (hSCRO) committee protocols. Informed consent was received for all participants.

## Data Availability

RNA sequencing data referenced in Figure 1-figure supplement 2 is available through Gene Expression Omnibus: GSE157652. Any additional data presented in this paper will be available from the authors upon request.

## Supplementary Figure Legends

**Figure 1-figure supplement 1:**
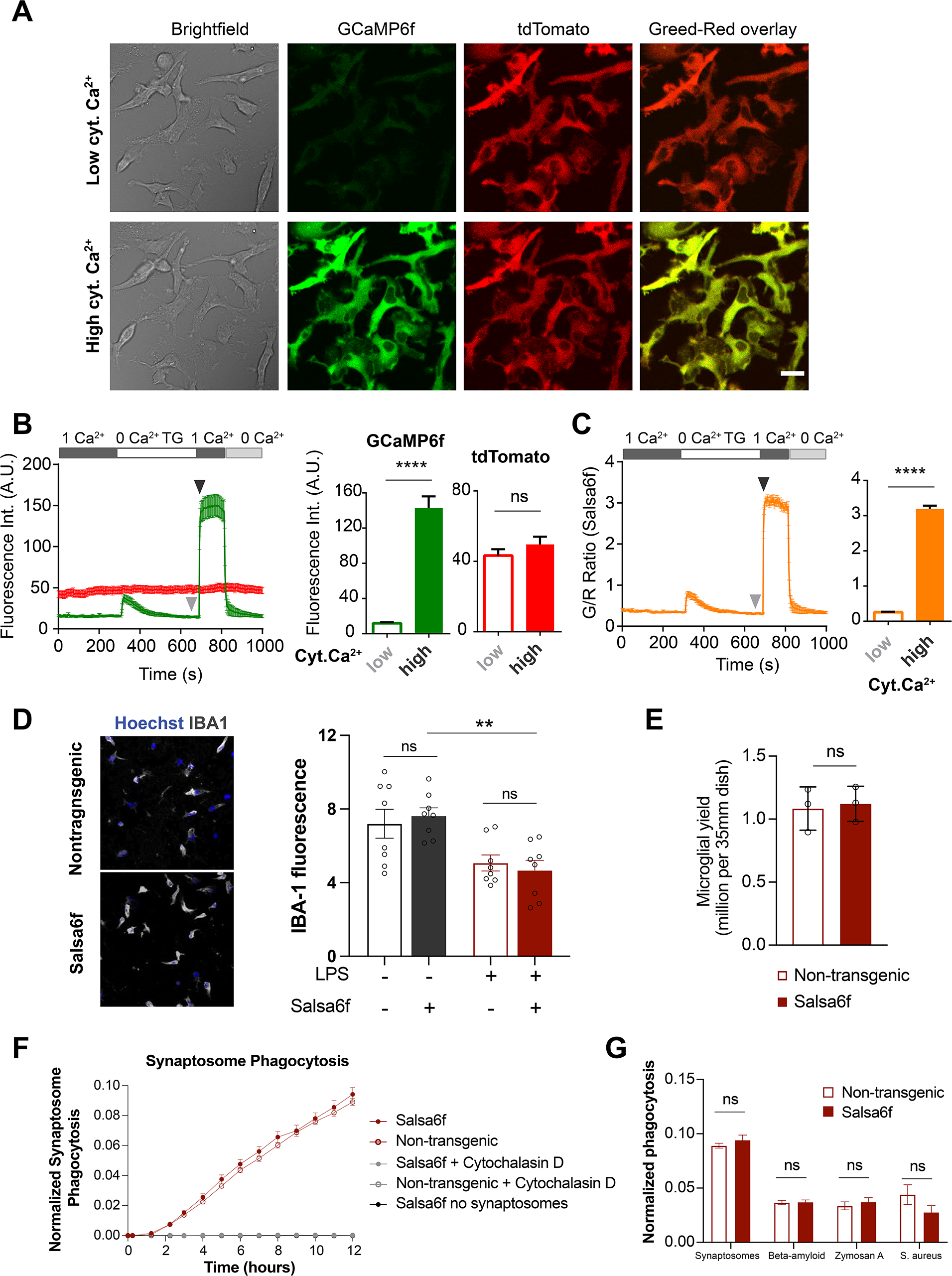
Validation of Salsa6f transgenic iPSC-microglia. (**A**) Representative bright field, green (GCaMP6f), red (tdTomato), and Green/Red channel overlay images of transgenic Salsa6f expressing iPSC-microglia at low (top row) and high (bottom row) cytosolic Ca^2+^ levels. Cells were treated with 2 μM thapsigargin (TG) to deplete stores and evoke store-operated Ca^2+^ entry (SOCE). Images are shown at the end of TG treatment for low Ca^2+^ and at the peak of SOCE for high Ca^2+^. Scale bar = 20 μm. (**B**) Trace of average change in fluorescence intensity of tdTomato (red) and GCamp6f (green) over time. Summary of GCaMP6f and tdTomato intensities before and after invoking SOCE are shown on the right. (**C**) Ratiometric GCaMP6f/ tdTomato signal (Green/ Red or G/R Ratio) over time calculated from (**B**). Summary of G/R Ratio at low and high cytosolic Ca^2+^ (**B-C**, n=19 cells, Mann-Whitney test). (**D**) Immunofluorescence images showing staining for the microglia-specific marker IBA1 in either resting or activated WT or Salsa6f-transgenic iPSC-microglia (left). Right panel shows quantification of IBA1 protein expression (n=4 wells, 2 independent images per well, t-test). Cells were activated with 100 ng/mL LPS (lipopolysaccharide for 24 hours). (**E**) Microglia cell counts at final day of differentiation (n=3 wells, t-test). (**F**) Phagocytosis of synaptosomes in WT non-transgenic (open circle) and Salsa6f-expressing (closed circle) iPSC-microglia. Cytochalasin D (grey, 10 µM) used as negative control to inhibit phagocytosis. Live cultures imaged on IncuCyte S3 (n=4 wells; 4 images per well). (**G**) Phagocytic load at 24 hr for synaptosomes, beta-amyloid, zymosan A, and *S. aureus* (n=4 wells; 4 images per well; one way ANOVA with Tukey post-hoc test). Data shown as mean ± SEM for traces and bar-graphs. *P values* indicated by **ns** for non-significant, *\**\*\*\* for *P < 0.0001*.

**Figure 1-figure supplement 2:**
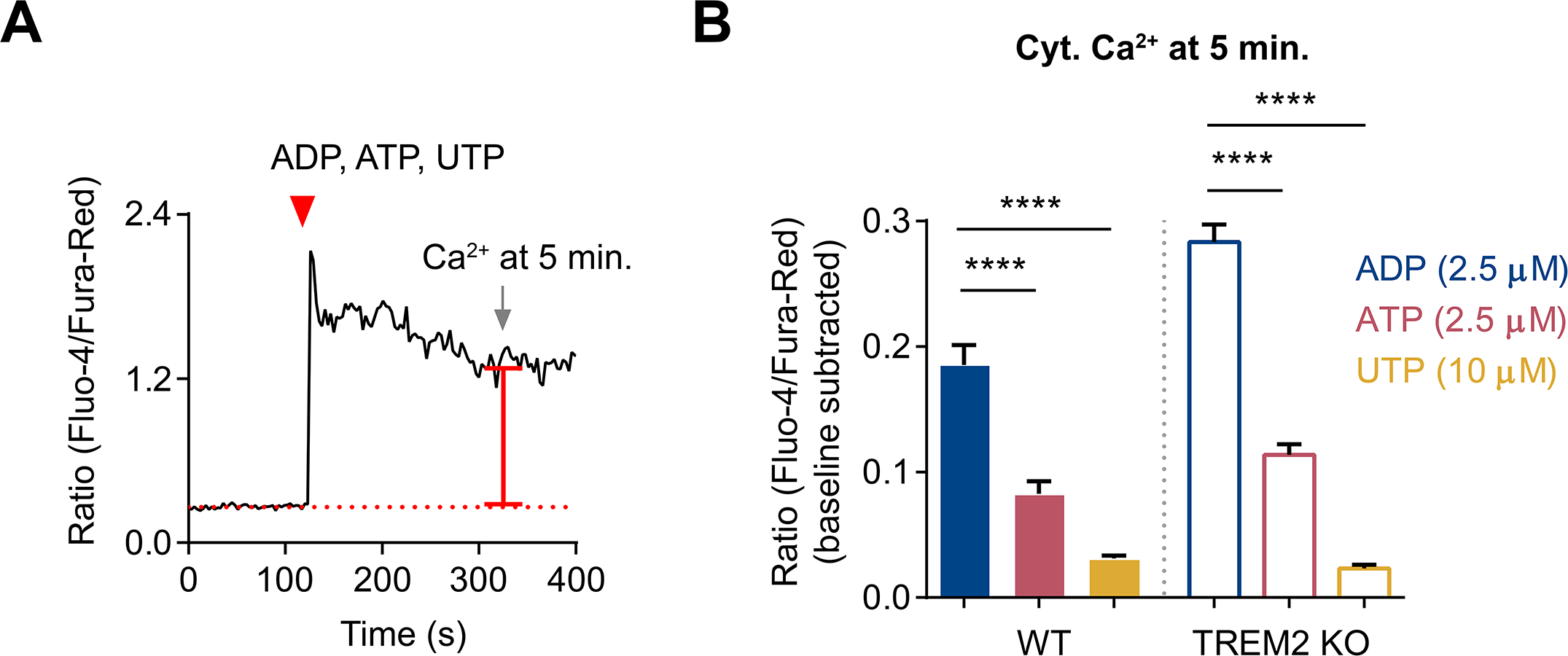
Comparison of cytosolic Ca^2+^ signal over time triggered by various purinergic agonists. (**A**) Representative trace showing changes in cytosolic Ca^2+^ in a single cell to illustrate the scheme for measuring cytosolic Ca^2+^ level 5 min after agonist application. (**B**) Bar-graph summary of cytosolic Ca^2+^ levels in WT and TREM2 KO iPSC-microglia 5 min after application of 2.5 μM ADP (blue), 2.5 μM ATP (red), and 10 μM UTP (yellow). N=165-274 cells pooled from 2-3 experiments. One-way ANOVA with multiple comparisons. Data shown as mean ± SEM for the bar-graph. *P values* indicated by *\**\*\*\* for *P < 0.0001*.

**Figure 2-figure supplement 1:**
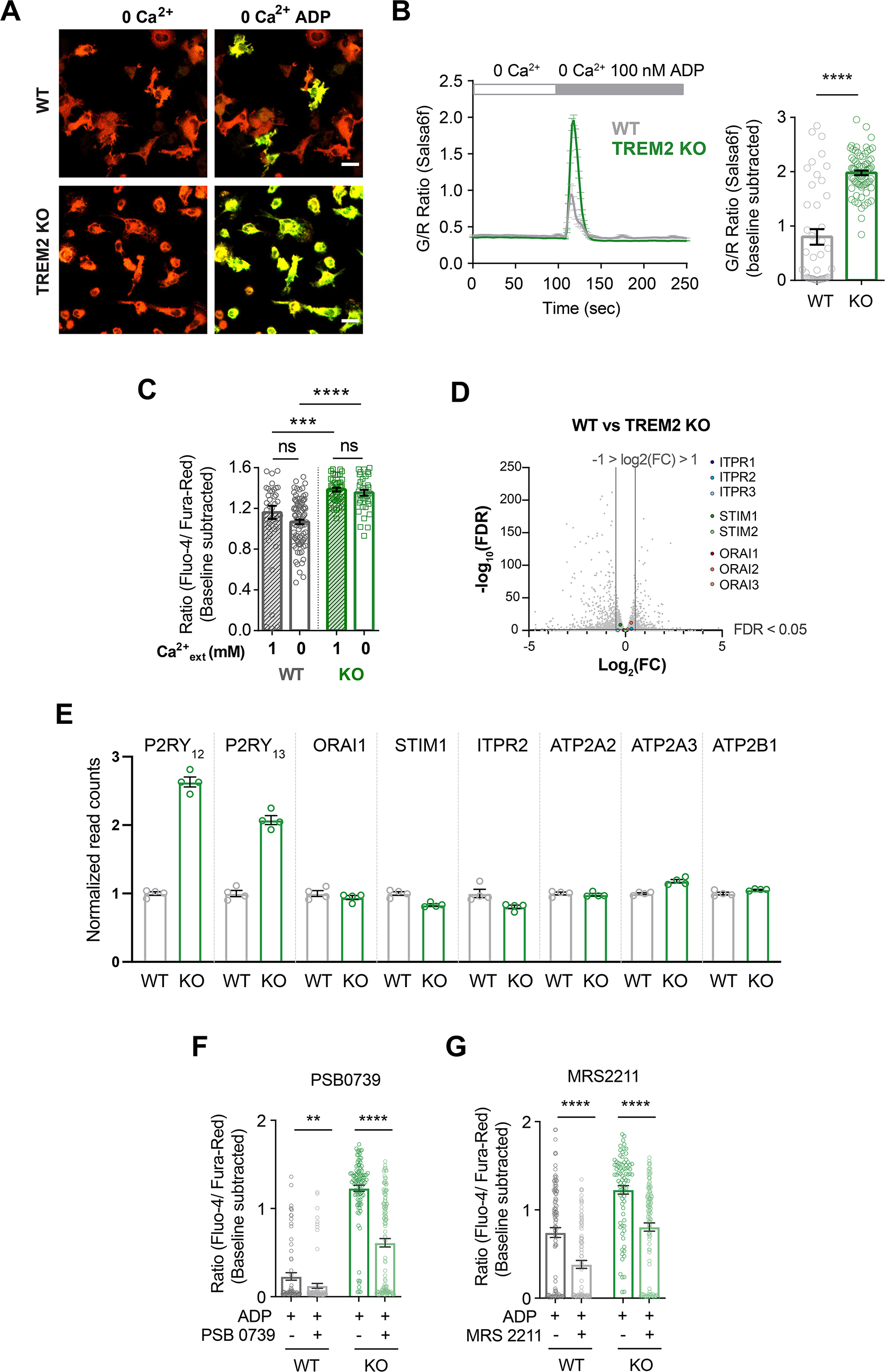
Role of P2Y_12_ and P2Y_13_ receptors in ADP-mediated augmentation of store-release in TREM2 KO microglia. (**A**) Representative green (GCaMP6f) and red (tdTomato) channel overlay images of WT (top) and TREM2 KO (bottom) iPSC-microglia before and peak Ca^2+^ response after ADP addition in Ca^2+^-free buffer. Scale bar = 20 mm. (**B**) Average trace (left panel) showing Ca^2+^ response to 100 nM ADP in Ca^2+^-free buffer. Quantification of peak signal (right panel, n=46-75 cells, 2 experiments, Mann-Whitney test). (**C**) Comparison of peak cytosolic Ca^2+^ in response to ADP (2.5 mM ADP) in 1 mM Ca^2+^ or Ca^2+^-free buffer (n=38-96 cells, ordinary one-way ANOVA with multiple comparisons). (**D**) Volcano plot of differentially expressed genes from bulk RNA-sequencing of WT and TREM2 KO iPSC-microglia (n=4). Genes for IP3R, STIM1, and ORAI1 are highlighted. (**E**) RNA normalized read counts for IP_3_ receptor type 2 (ITPR2), PMCA1 (ATP2B1), SERCA2 (ATP2A2), SERCA3 (ATP2A3), STIM1, and ORAI1 in WT and TREM2 KO iPSC-microglia. Isoforms expressed lower than 10 reads in any sample are not considered expressed and are not shown. Relative expression of P2Y_12_ and P2Y_13_ receptors are shown for comparison of the relative fold-change between WT and TREM2 KO cells. (**F-G**) Peak Ca^2+^ response in Ca^2+^ free buffer after treatment with 1 or 10 μM ADP in the presence of P2Y_12_ receptor antagonist PSB 0739 (**F**) or P2Y_13_ receptor antagonist MRS 2211 (**G**), respectively. Cells were pretreated with 10 μM of PSB 0739 or 10 μM MRS 2211 for 30 min before imaging. (72-128 cells, **F**; 83-117 cells, **G**; representative of 3 experiments, Mann-Whitney Test). Data shown as mean ± SEM for traces and bar-graphs. Data shown as mean ± SEM for traces and bar-graphs. *P values* indicated by **ns** for non-significant, *\**\* for *P < 0.01, \**\*\* for *P < 0.001, \**\*\*\* for *P < 0.0001*.

**Figure 3-figure supplement 1:**
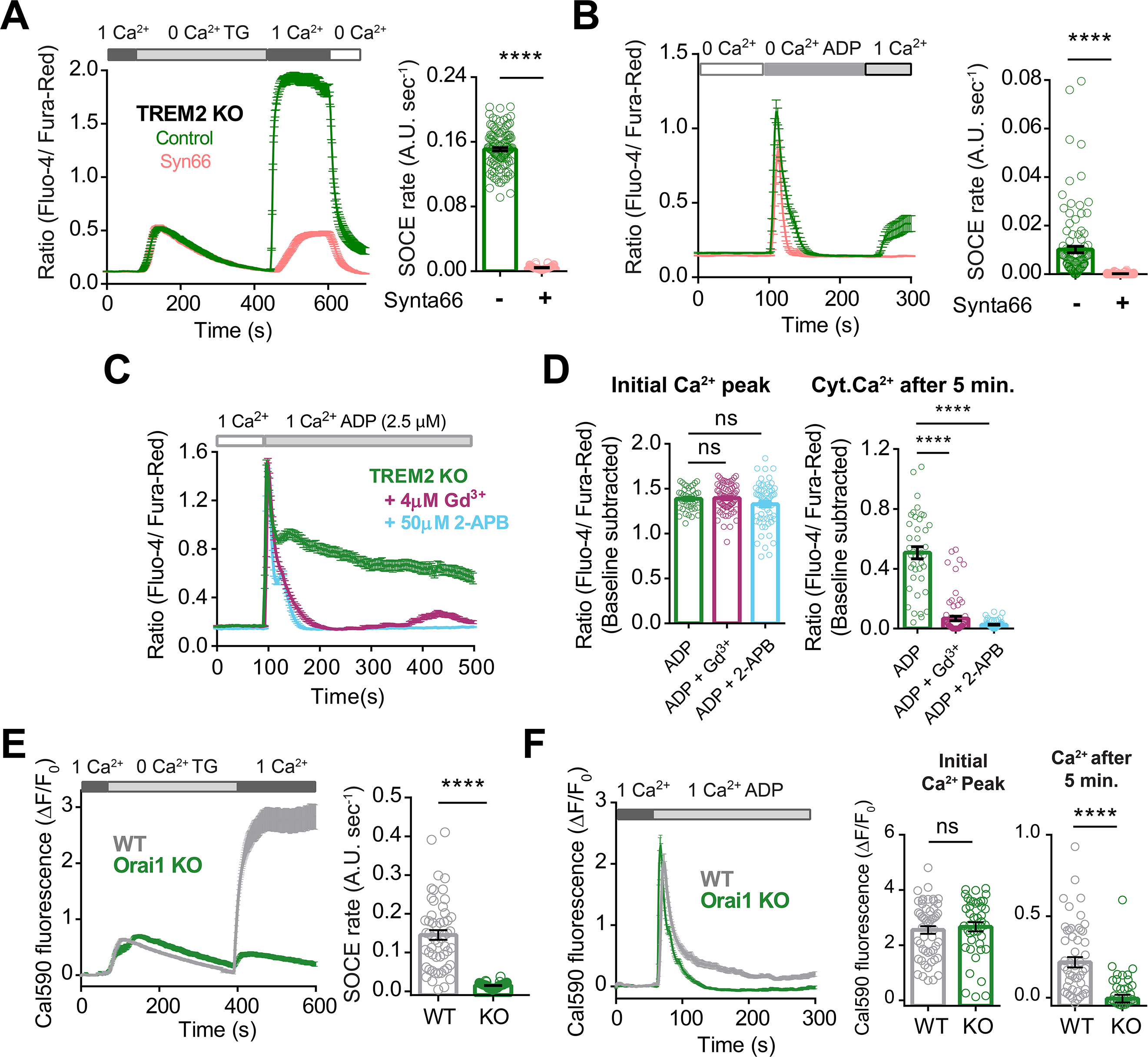
Regulation of SOCE in iPSC-microglia. (**A**) Average trace showing SOCE triggered in TREM2 KO microglia via emptying ER Ca^2+^ stores with thapsigargin (TG, 2 μM) in Ca^2+^-free buffer followed by re-addition of 1 mM Ca^2+^ in the absence (control, green trace) or presence (red trace) of the Orai channel inhibitor Synta66. Cells were pretreated with Synta66 (10 μM) for 30 min before experiment. Bar-graph summary of the rate of Ca^2+^ influx after re-addition of 1 mM Ca^2+^ (80-126 cells, Mann-Whitney test). (**B**) SOCE evoked by ADP (2.5 μM) in TREM2 KO microglia (green trace), using a similar Ca^2+^ addback protocol. Red trace shows effect of Synta66 on ADP-evoked SOCE. Right panel summarizes the rate of ADP-triggered Ca^2+^ influx after re-addition of 1 mM Ca^2+^ (n= 125-154 cells, 2 experiments, Mann-Whitney test). (**C-D**) Cytosolic Ca^2+^ response to ADP in TREM2 KO iPSC-microglia pre-treated with 2-APB (50 μM) or Gd^3+^ (5 μM) to block store-operated Ca^2+^ entry (SOCE). Average traces (**C**), baseline-subtracted initial peak Ca^2+^ responses to ADP (**D**, left panel), and baseline-subtracted Ca^2+^ after 5 min of ADP addition (**D**, right panel) are shown (n= 41-74 cells, ordinary one-way ANOVA with multiple comparisons). (**E-F**) Role of Orai1 in TG- and ADP-evoked SOCE in iPSC-microglia. (**E**) Comparison of TG-evoked SOCE in WT and Orai1 KO cell showing average traces (left panel) and summary of SOCE rate (right panel; n= 42-54 cells, 3-4 experiments, Mann-Whitney test). (**F**) ADP-evoked SOCE in WT and Orai1 KO showing average traces (left panel) and summary of SOCE rate (right panel; n= 42-53 cells, 3-4 experiments, Mann-Whitney test). Data shown as mean ± SEM for traces and bar-graphs. *P values* indicated by **ns** for non-significant, *\**\*\*\* for *P < 0.0001*.

**Figure 3-figure supplement 2:**
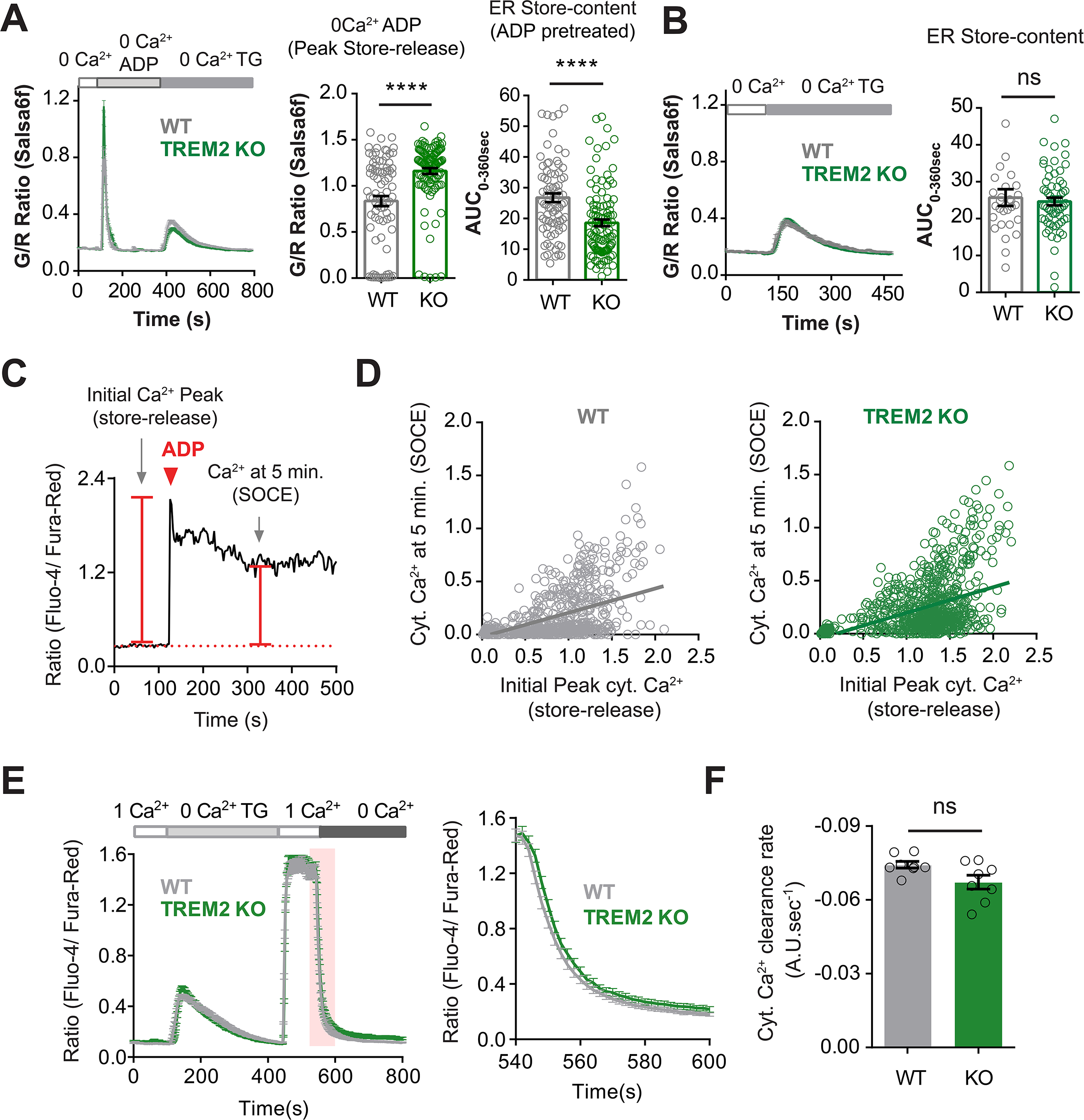
ADP depletes ER Ca^2+^ stores to a greater extent in TREM2 KO microglia. (**A**) TG-pulse experiment to measure residual ER Ca^2+^ pool in cells after initial treatment with ADP (1 μM) and subsequent treatment with thapsigargin (2 μM). Imaging was done in Ca^2+^-free buffer to prevent Ca^2+^ influx across the PM. Average trace (left panel), Peak ADP Ca^2+^ response (middle panel) and extent of TG-induced ER store-release measured as area under curve (AUC, right panel) (n=81-108 cells, Mann-Whitney test). (**B**) Control experiment comparing the ER-Ca^2+^ pool in WT and TREM2 KO microglia after store-depletion with TG, and without any pretreatment with ADP (n=29-63 cells, Mann-Whitney test). (**C-D**) Relationship between ADP-induced store-release and SOCE in iPSC-microglia. (**C**) Representative single-cell trace of Ca^2+^ signal in response to ADP in 1 mM extracellular Ca^2+^ buffer showing the scheme for measuring ER store-release as the initial Ca^2+^ peak and SOCE as cytosolic Ca^2+^ level 5 min after ADP application. (**D**) Scatter-plot showing correlation of initial ADP-induced Ca^2+^ response (store-release) and cytoplasmic Ca^2+^ after 5 min (SOCE) in WT (grey) and KO (green) cells (n=866-935 cells from multiple imaging runs with a range of ADP doses; in μM: 0.001, 0.1, 0.5, 1, 2, 2.5, 5, 10. Comparison of slopes between WT and TREM2 KO: *P = 0.7631*; Extra sum of squares F test). (**E-F**) Comparison of cytosolic Ca^2+^ clearance indicative of PMCA pump activity in WT and TREM2 KO microglia. SOCE was invoked and rate of Ca^2+^ decline was measured after addition of 0 mM Ca^2+^. (**E**) Average trace showing invoking SOCE with 2 μM TG (left panel). Right panel shows the drop in cytosolic Ca^2+^ following addition of Ca^2+^ free solution as highlighted (pink) in the SOCE trace (**F**) Summary of rate of Ca^2+^ decline after addition of 0 mM Ca^2+^ (n= 8 imaging fields, 142-175 total cells, Mann-Whitney Test). Data shown as mean ± SEM for traces and bar-graphs. *P values* indicated by **ns** for non-significant, *\**\* for *P < 0.01*.

**Figure 4-figure supplement 1:**
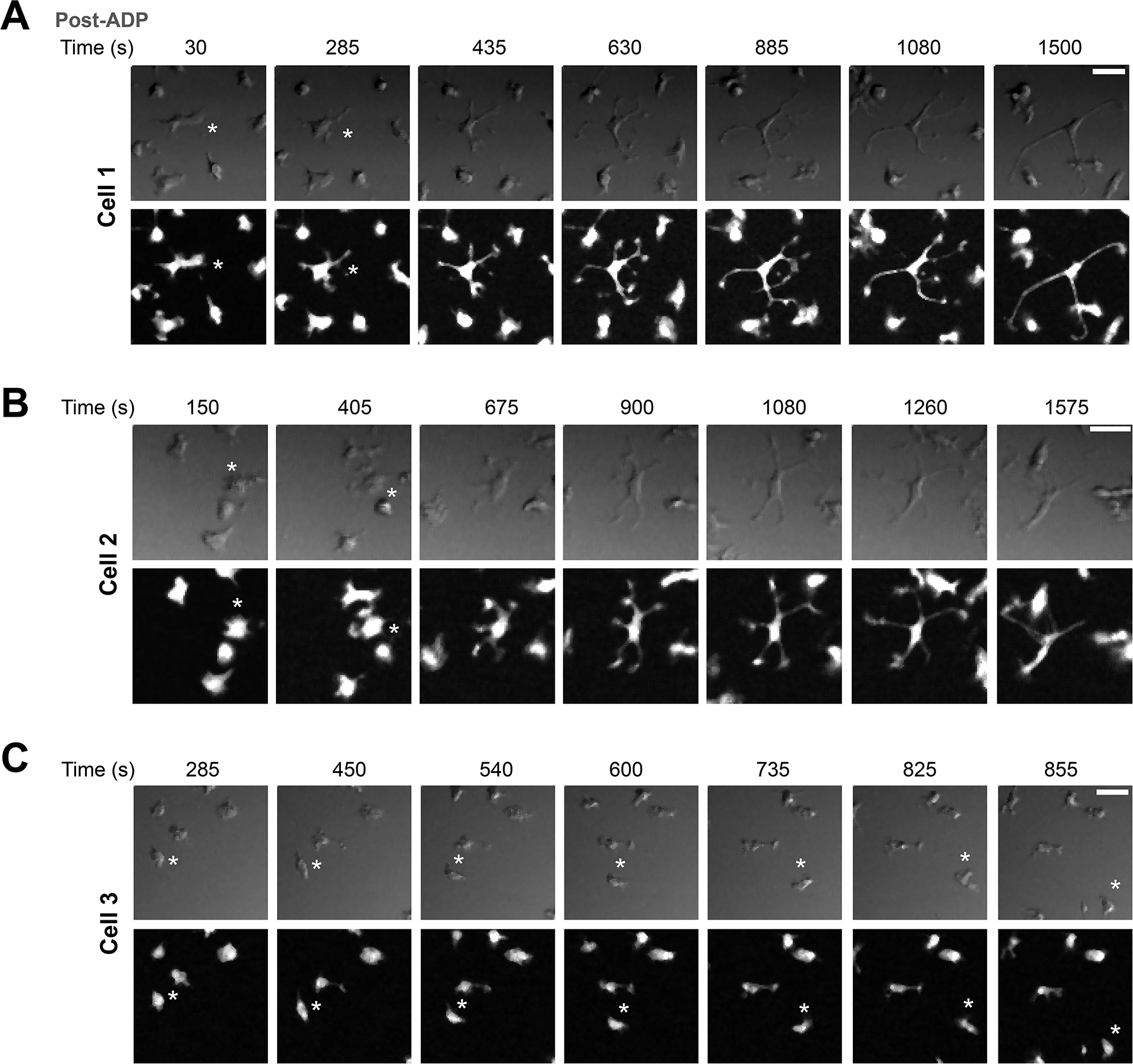
ADP-mediated process extension in WT iPSC-microglia. **(A)** Representative images of a cell (Cell 1) from a time-lapse experiment showing increased branching and extension of processes in GFP-expressing WT iPSC-microglia, at times indicated following addition of 2.5 μM ADP. Bright field DIC images (top row) and GFP images (bottom row) are shown. (**B**) Another example of a cell (Cell 2) showing process extension in the same imaging field. (**C**) A motile cell (Cell 3) in the same imaging field is shown for comparison. Note the lack of displacement in cells that extend their process, and lack of significant process extension in a highly motile cell. Scale bar: 15 μM.

**Figure 5-figure supplement 1:**
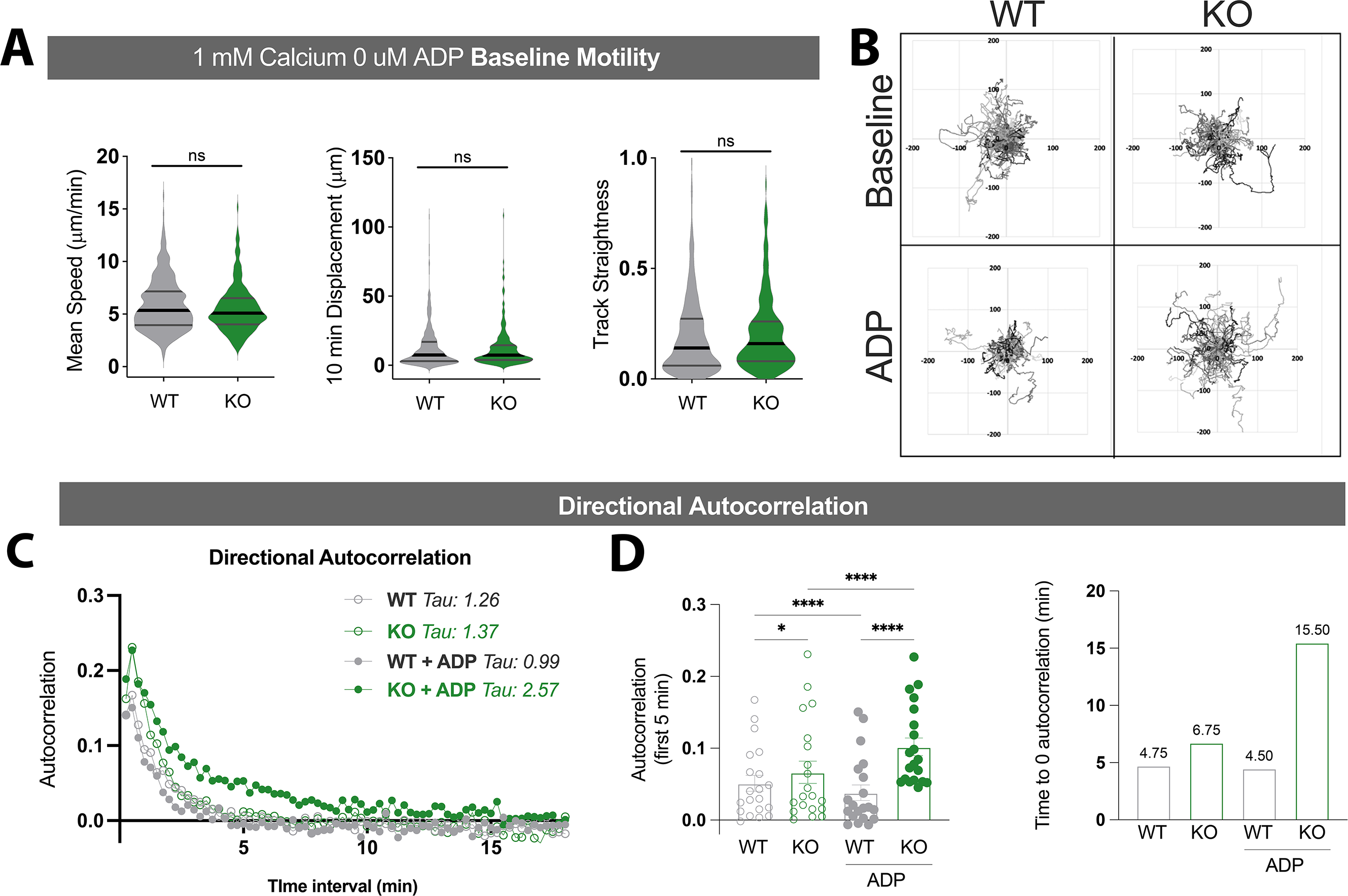
Motility analysis in WT and TREM2 KO iPSC-microglia. (**A**) Summary of microglial mean speeds, displacement over 10 min, and track straightness in open-field migration in the absence of any purinergic stimulation (student’s t-test). (**B**) Flower plots show similar displacement from origin for WT (left) and TREM2 KO (right) cells. (**C**) Directional Autocorrelation calculated via DiPer excel macro. Due to lack of directional gradient, directional autocorrelation of motility vectors is expected to drop quickly. Time constants for best-fit single exponential curves are indicated, consistent with increased straightness for TREM2 KO cells treated with ADP. (**D**) Directional autocorrelation of WT (grey) and TREM2 KO (green) iPSC-microglia at baseline (open circles) or after ADP addition (filled circles). Mean autocorrelation values in the first 5 min (left panel, one-way ANOVA) and time (min) until autocorrelation reaches zero (right panel). Data shown as mean ± SEM for the bar-graph in (D), and as violin plots with mean, 25^th^ and 75^th^ percentile in (A). *P values* indicated by **ns** for non-significant, * for *P < 0.05,* and **** for *P < 0.0001*.

**Figure 5-figure supplement 2:**
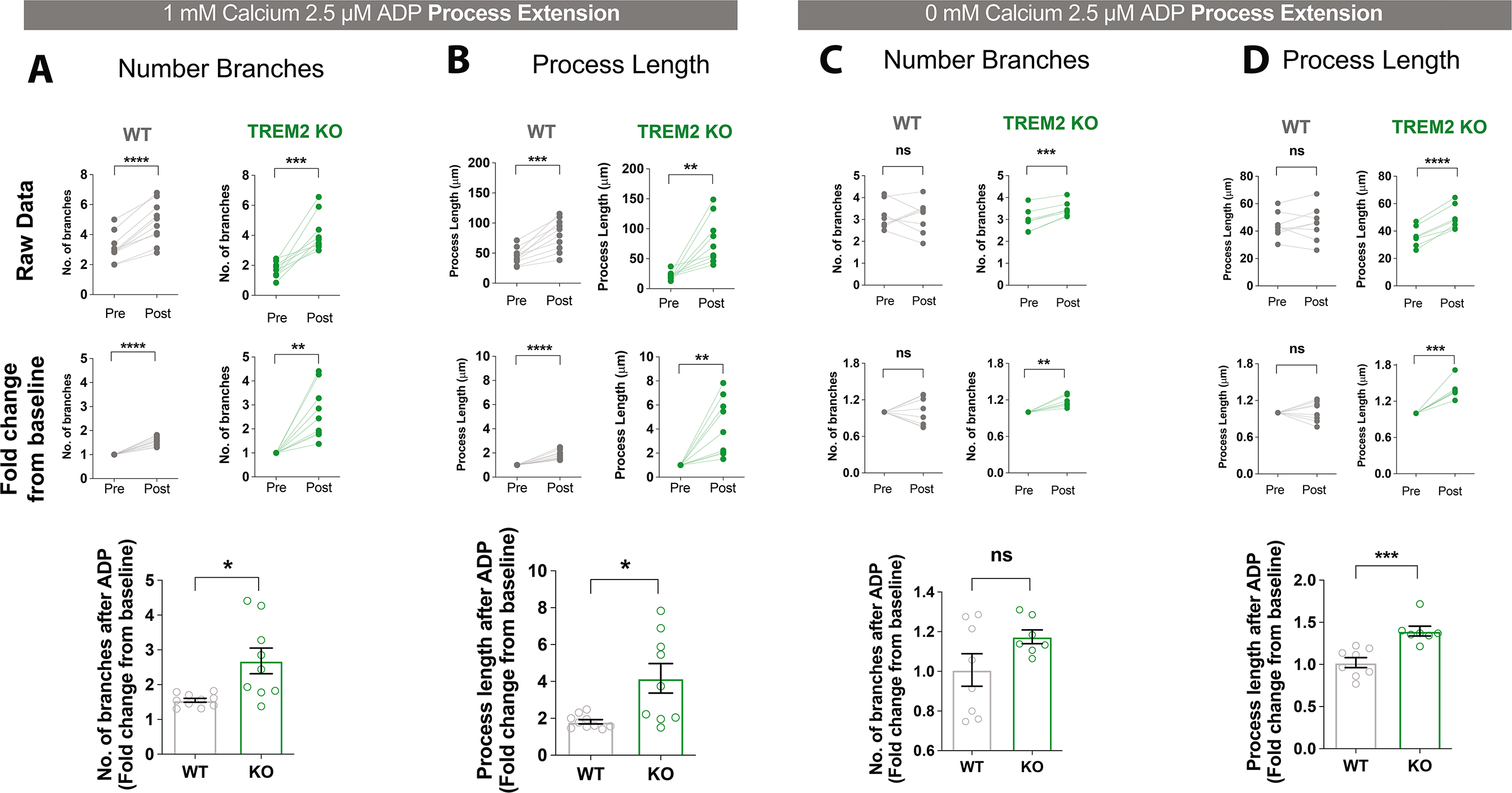
Comparison of process extension in WT and TREM2 KO Microglia. Branching and process extension in WT and TREM2 KO iPSC-microglia 30 min after addition of ADP in 1 mM (**A, B**) or 0 mM extracellular Ca^2+^ buffer (**C, D**). (**A**) Data displayed as paired-plots showing average branch number per cell in an imaging field (top row) and normalized to pre-ADP values for each imaging field (middle row). Bottom row shows fold change in branching after ADP treatment for WT (grey) and KO (green) iPSC-microglia. (**B**) Changes in process length in the same dataset as **A**. n=151-158 cells, WT; 133-167 cells, KO; 9-10 imaging fields, 3-4 experiments. (**C, D**) Same analysis as **A, B** but with ADP in Ca^2+^ free buffer. n=137-143 cells, 8 imaging fields, 2-3 experiments. (**A-D**) p-values calculated by two-tailed paired Students t-test for the paired-plots, and by unpaired t-test when comparing fold-change in WT and KO cells. Data shown as paired plots and as mean ± SEM for the bar-graphs. *P values* indicated by **ns** for non-significant, * for *P < 0.05, \**\* for *P < 0.01, \**\*\* for *P < 0.001* and **** for *P < 0.0001*.

**Figure 6-figure supplement 1:**
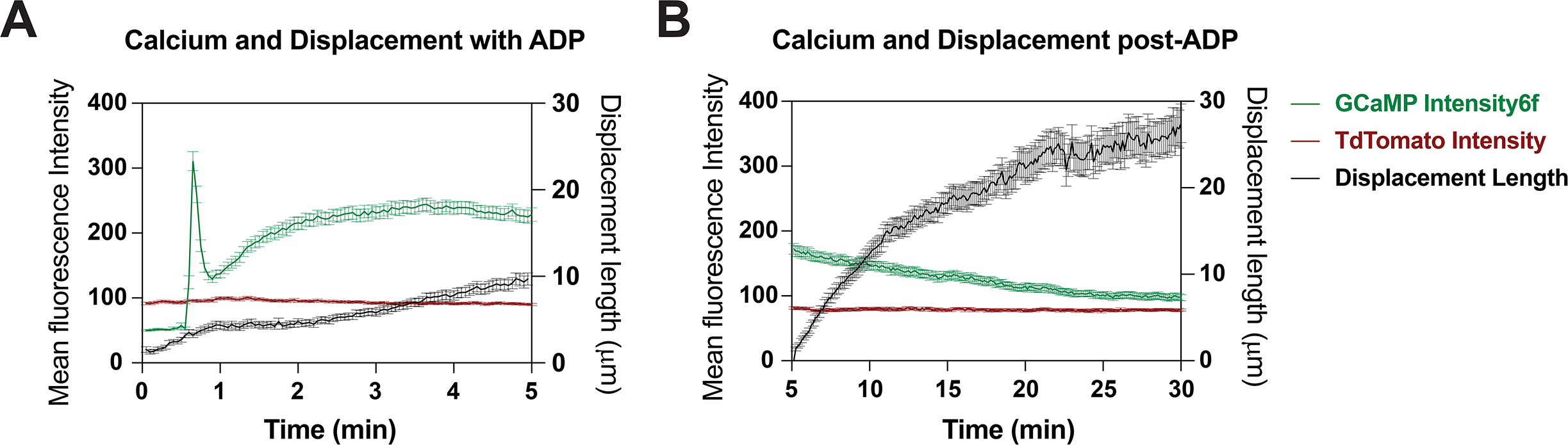
Tracking cell motility and cytosolic Ca^2+^ using Salsa6f-expressing iPSC cell-line. (**A**) Average change in single cell fluorescence intensity of tdTomato (red trace) and GCaMP6f (green trace) (left Y-axis) in WT Salsa6f microglia over 5 min following ADP treatment, overlaid with corresponding change is cell displacement over time (black trace, right Y-axis) (n=52-79 cells). (**B**) Same as (A) but for cells tracked over a period of 30 min. Data shown as mean ± SEM for average traces.

**Figure 6-figure supplement 2:**
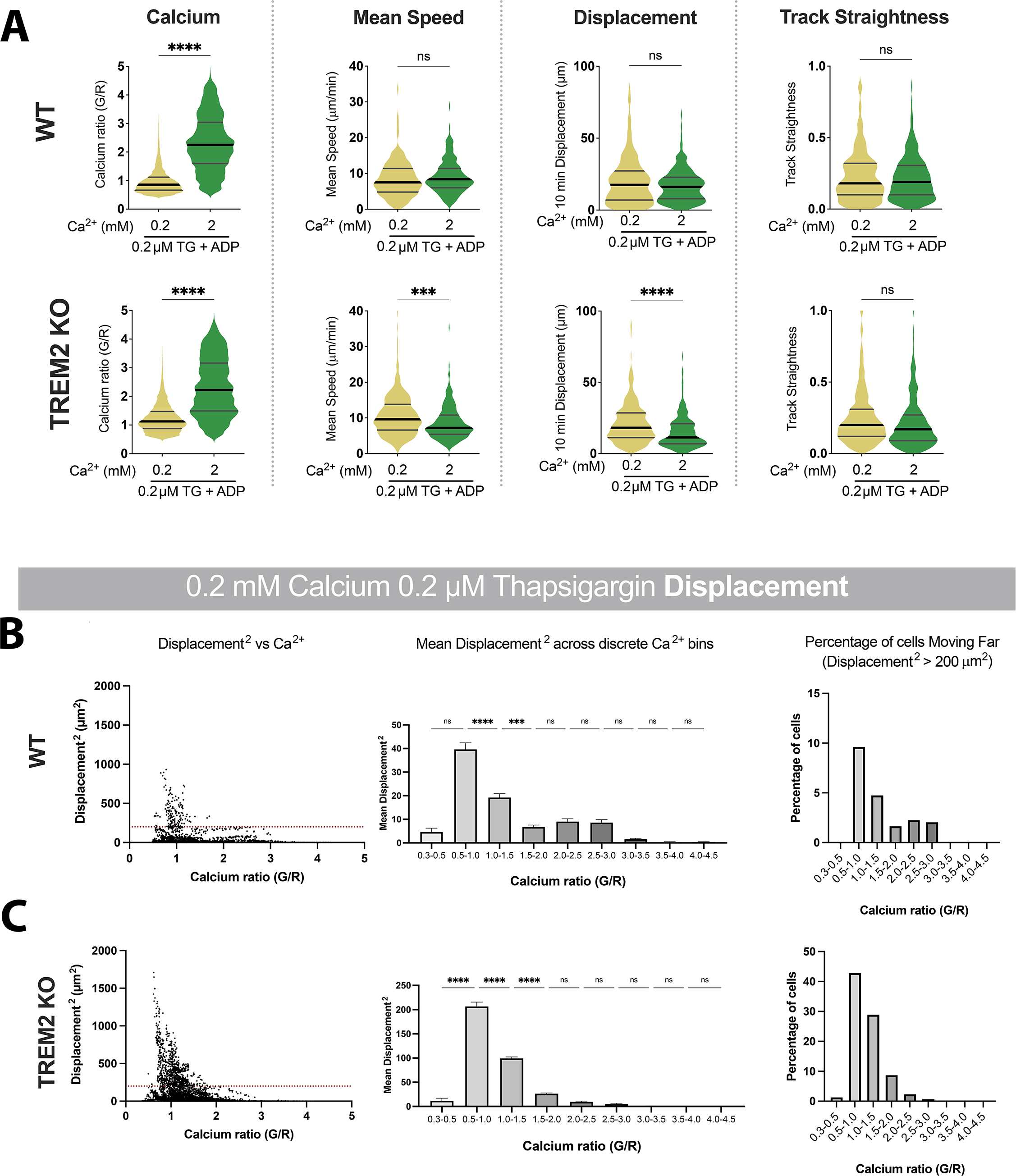
Motility analysis with varying Ca^2+^. (**A**) Salsa6f Ca^2+^ ratios and microglia motility in WT (top) and KO (bottom) microglia, with ADP added: yellow (0.2 mM Ca^2+^, TG + ADP), green (2 mM Ca^2+^, TG + ADP). Cytosolic Ca^2+^ levels indicated by instantaneous single-cell G/R Ratio. Mean of instantaneous speeds, 10 min track displacement and track straightness calculated as before. Students t-test **** p < 0.0001; *** p = 0.0001. n=164-393 cells. (**B, C**) Ca^2+^ dependence of track displacement length in 0.2 mM Ca^2+^ in WT cells (**B**) and TREM2 KO cells (**C**). Correlation between instantaneous Ca^2+^ and frame-to-frame displacement (left panels). Each dot represents an individual cell for an individual frame. Dotted red line represents displacement of 200 μm^2^. Mean square of frame-to-frame displacement of cells binned by instantaneous G/R Ca^2+^ ratio (middle panels, 1-way ANOVA **** p < 0.0001). Each data point is calculated for a bin increment of 0.5 G/R ratio. Summary of cells with frame-to-frame square displacement > 200 μm^2^ (right panels). WT cells (**B**) displace less than KO cells (**C**). For each cell type, larger displacements are correlated with lower G/R Ca^2+^ ratios. Cells which maintain elevated cytoplasmic Ca^2+^ do not displace as far. For WT: p < 0.0001; r = −0.4778; number pairs = 5973. For KO: p < 0.0001; r = −0.3699; number pairs = 5761 (Spearman’s correlation). Data shown as mean ± SEM for bar-graphs (B, C) and as violin plots with mean, 25^th^ and 75^th^ percentile (A). *P values* indicated by **ns** for non-significant, *\**\*\* for *P < 0.001* and **** for *P < 0.0001*.

**Figure 7-figure supplement 1:**
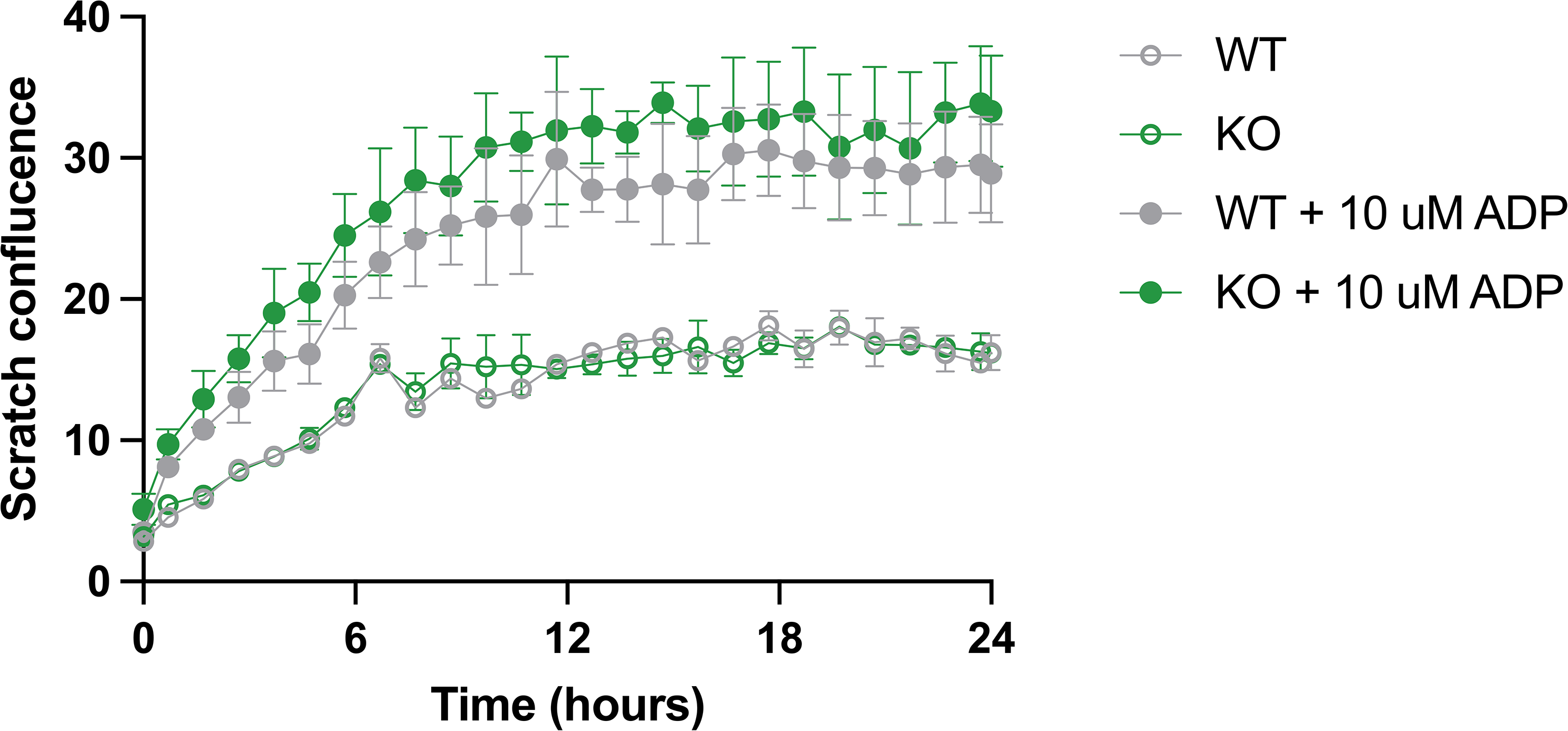
TREM2 WT and KO close scratch wound at similar rates. Scratch closure over 24 hours in WT (grey) and TREM2 KO (green) iPS-microglia with (filled symbols) or without (empty symbols) pre-stimulation of iPSC-microglia with ADP (10 μM, 30 min). N=2 wells, 2 images per well. Data shown as mean ± SEM.

